# LC-MS/MS-PRM Quantification of IgG glycoforms using stable isotope labeled IgG1 Fc glycopeptide standard

**DOI:** 10.1101/2022.08.02.501850

**Authors:** Miloslav Sanda, Qiang Yang, Guanghui Zong, He Chen, Zhihao Zheng, Harmeet Dhani, Khalid Khan, Alexander Kroemer, Lai-Xi Wang, Radoslav Goldman

## Abstract

Targeted quantification of proteins is a standard methodology with broad utility, but targeted quantification of glycoproteins has not reached its full potential. The lack of optimized workflows and isotopically labeled standards limits the acceptance of glycoproteomics quantification. In this paper, we introduce an efficient and streamlined chemoenzymatic synthesis of a library of isotopically labeled glycopeptides of IgG1 which we use for quantification in an energy optimized LC-MS/MS-PRM workflow. Incorporation of the stable isotope labeled N-acetylglucosamine enables an efficient monitoring of all major fragment ions of the glycopeptides generated under the soft collision induced dissociation (CID) conditions which reduces the CVs of the quantification to 0.7-2.8%. Our results document, for the first time, that the workflow using a combination of stable isotope labeled standards with intra-scan normalization enables quantification of the glycopeptides by an electron transfer dissociation (ETD) workflow as well as the CID workflow with the highest sensitivity compared to traditional workflows., This was exemplified by a rapid quantification (13-minute) of IgG1 Fc glycoforms from COVID-19 patients.

**Graphic Abstract:** 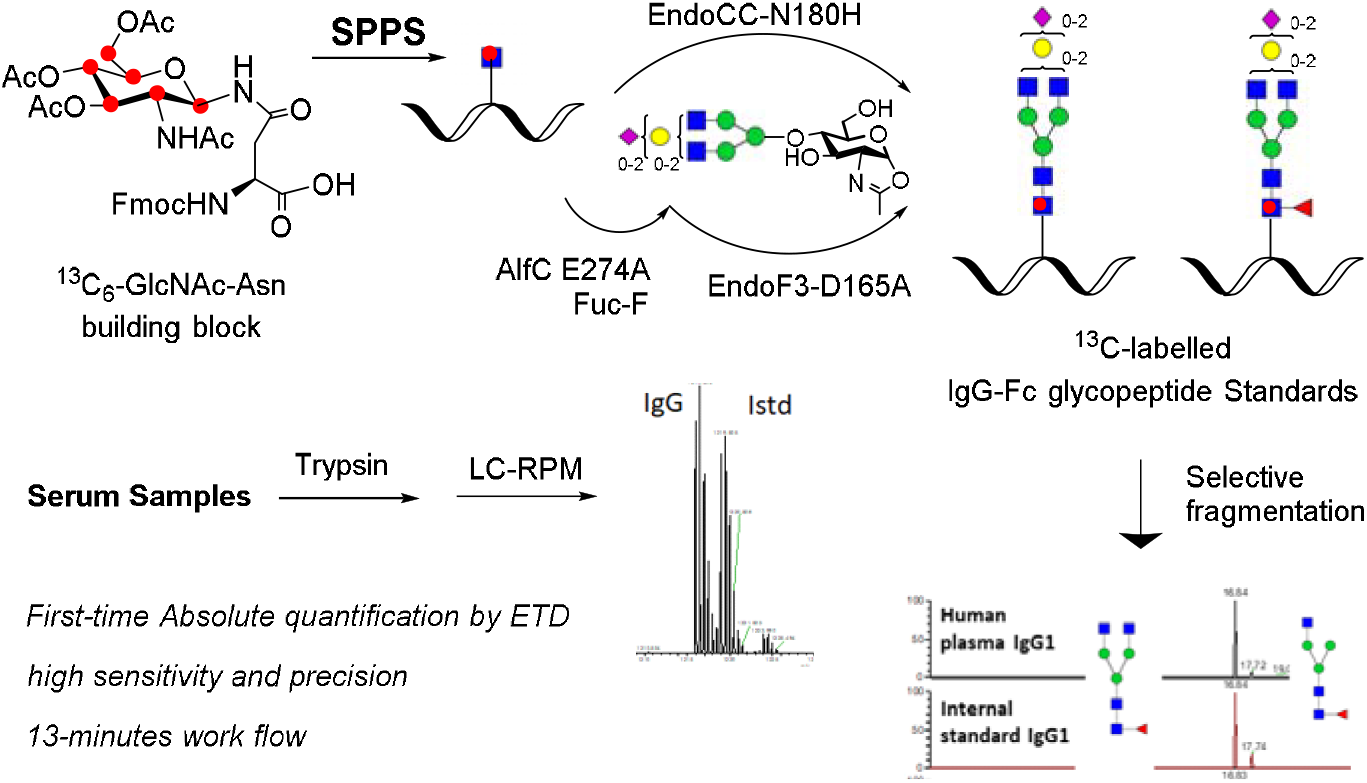

## INTRODUCTION

N-glycosylation is a common and complex post-translational modification of proteins ^1–3^ whose impact on an organism increases with its complexity^4^. Defects in this pathway in humans lead to congenital disorders of glycosylation (CDG) often incompatible with life ^5^. The N-glycosylation process is initiated in the endoplasmic reticulum by an oligosaccharyl transferase (OST) complex ^6^ which transfers a common lipid-linked N-glycan building block to an asparagine in an NXS/T sequon (X≠P)^7^. The attached N-glycans are trimmed and further expanded by multiple glycosidases and glycosyltransferases during the maturation of the secreted/membrane proteins in the Golgi apparatus. Appropriate glycosylation is critical in many important molecular recognition processes, including protein folding, protein trafficking, and protein-protein/glycan interactions^8–10^. Perhaps the most studied glycoprotein is the immunoglobulin G (IgG) ^11, 12^. Glycosylation of the N297 of human IgG is known to modulate interactions with the Fc receptors ^13^ and subsequent biological^14^ and therapeutic^15^ responses. It is therefore of considerable interest to quantify accurately the IgG glycoforms. So far, many analytical methods have been introduced for this purpose ^16–23^, including mass spectrometric methods for relative quantification of the IgG glycopeptides ^22, 23^. In spite of these advances, the quantification of IgG glycoforms and other glycopeptides by targeted mass spectrometric methods has been limited,^21^ in contrast to the quantification of metabolites, drugs or proteins, for which well-established analytical approaches with clinical utility have been established ^25–27^. One reason for the limited acceptance of IgG glycoform quantification in clinical samples is the dominant production of less specific glycan fragments (oxonium ions) under CID conditions used typically for the fragmentation of glycopeptides ^28–30^. Another reason is the lack of synthetic isotope labeled standards (SIS) of glycopeptides which present a substantial synthetic challenge despite recent advances in the chemical and chemoenzymatic synthetic approaches ^31–34^. In the course of our study, Li and coworkers have reported a chemoenzymatic synthesis of an array of fucose isotope-labeled Fc glycopeptides for their absolute quantitation but the method is limited to core-fucosylated glycopeptides ^35^. In this study, we report a more efficient modular and streamlined synthetic route for both core-fucosylated and non-fucosylated glycopeptides ^35^. Accordingly, we advance the quantification of the IgG glycoforms by introduction of the SIS glycopeptides in combination with our energy optimized targeted mass spectrometric quantification workflow.

## MATERIALS and METHODS

### Synthesis of stable isotope-labeled IgG1 Fc glycopeptides

D-[UL-^13^C6]-N-Acetylglucosamine was purchased from Cambridge Isotope Lab, Inc. Fmoc-amino acids were purchased from ChemPep, Inc. All other chemicals, reagents, and solvents were purchased from Sigma–Aldrich.

### SPPS of ^13^C-labeled IgG1-Fc-GlcNAc peptide (IgG1-FcP-Gn, compound 5)

Preparation of C-labeled IgG1-FcP-Gn acceptor was performed under microwave synthesis conditions using a CEM Liberty Blue microwave peptide synthesizer. Synthesis was based on Fmoc chemistry using Rink Amide resin (0.66 mmol/g) on a 0.1 mmol scale, following the protocol as described by Zong et al, with incorporation of C-labeled GlcNAc-Asn. The crude peptides were purified by RP-HPLC and the purity and identity were confirmed by analytical HPLC and LC-MS analysis. An unlabeled identical peptide was synthesized using the same protocol as well.

### Synthesis of ^13^C-labeled IgG1-Fc-GlcNAcFuc peptide (IgG1-FcP-GnF, compound 5)

^13^C-labeled IgG1-FcP-GnF was synthesized by transfer of fucosyl moiety to ^13^C labeled IgG1-FcP-Gn with a fucoligase AlfC-E274A, using α-L-Fucosyl fluoride as the donor, following published procedures ^36, 45^.

### Preparation of various glycan oxazolines

Various biantennary complex glycans were prepared from a combination of mild acid treatment and enzyme trimmings from sialylglycopeptide (SGP) that was isolated from egg yolk powder^43^. First, the SGP was partially desialyted with trifluoro acid (TFA). In a typical protocol, 220 mg SGP was treated with 0.4% of trifluoro acid TFA (pH ~2) at 50 °C for 2-4 h to reach approximately 50% desialylation. The partially desialylated SGP mixture was neutralized with 1 M NaOH and then cleaved with an endoglycosidase Endo-S2 ^42^ to dissociate the glycan and peptide portions. After desalting with a Sephadex G-10 column, the S1G2, S2G2, and G2 glycan were separated with anion exchange chromatography on a HiTap Q-XL column with a 0 to 0.2 M NaCl gradient. G2 glycan that was mixed with peptide portion was further purified with cation exchange with a HiTrap Q-XL column in a pass-through mode. G0 glycan was obtained by the treatment of G2 glycan with a ß1,4-glactosidase, BgaA ^47^. To prepare G1 glycan, S1G2 glycan was trimmed with BgaA to generate S1G1 glycan, which was further processed with a neuraminidase, MvNA^40^ to afford targeted G1 glycan. All the glycans were converted to activated oxazolines for glycosidase-mutant catalyzed transglycosylation, following the previously reported procedure ^48^.

### Synthesis of C-labeled IgG1-Fc glycopeptides

Fucosylated glycopeptides were synthesized by the EndoF3 glycosynthase mutant EndoF3-D165A catalyzed transglycosylation, following our previously published procedure^38^. The product was purified by prep HPLC with a semiPrep HPLC column. None-fucosylated glycopeptides were synthesized by glycan transfer with the EndoCC mutant, EndoCC-N180H ^37^. In a typical EndoCC-N180H catalyzed reaction, 1 mg of ^13^C-labeled IgG1-Fc-GlcNAcFuc peptide was mixed with 3 mol. eq. of glycan oxazolines, 0.1 μg/μL the glycosynthase (EndoCC-N180H) in a phosphate buffer (100 μL, 50 mM, pH 7). The reaction mixture was incubated at 30 □ for 30-60 min, with LC-MS monitoring of reaction progression. 90-95% of glycan transfer were achieved under such condition. The final product was purified by prep HPLC. After lyophilization, the synthesized glycopeptides were weighed on an accurate balance and further quantitated by analytic HPLC with IgG1-FcP-GlcNAc as the internal standard.

### Patient enrollment and blood sample processing

Participants who were diagnosed with COVID-19 between March and July, 2020, using reverse transcriptase polymerase chain reaction for SARS-CoV-2, were enrolled in collaboration with the MedStar Georgetown Transplant Institute, MedStar Georgetown University Hospital and the Center for Translational Transplant Medicine, Georgetown University Medical Center, Washington, D.C. (**Supplemental table 2**), under protocols approved by the Georgetown University Medical Center IRB (Approval # STUDY00002359; IRB # 2017-0365). Samples obtained from participants before the COVID-19 era were used as controls. All participants provided written informed consent. Blood was collected in serum vacutainer (BD Vacutainer CPT; BD Biosciences) and processed within 12 hours of blood draw by centrifuging at 1200×g for 10 minutes. Aliquots of 0.5mL were placed into vials and stored at −80°C until further use. Aliquots of thawed serum were diluted 1:69 with a sodium bicarbonate solution and processed as described previously ^20^ with minor modifications. Briefly, diluted serum was reduced with 5 mM DTT for 60 min at 60°C and alkylated with 15 mM iodoacetamide for 30 min in the dark. Trypsin Gold (Promega, Madison, WI) digestion (2.5 ng/μl) at 37°C in Barocycler NEP2320 (Pressure BioSciences, South Easton, MA) for 1 hour, samples were evaporated using a vacuum concentrator (Labconco), and dissolved in mobile phase A (2% ACN, 0.1% FA). Tryptic peptides were analyzed without further processing to ensure reliable quantification of the glycoforms.

### Glycopeptide analysis by a nano LC-MS/MS-PRM workflow

Glycopeptide separation was achieved on an Ultimate 3000 nanochormatography system using a capillary analytical 75 μm x 150 PEPmap300, 3 μm, 300 Å column (Thermo) interfaced with an Orbitrap Fusion Lumos (Thermo). Glycopeptides were separated at 0.3 μl/min as follows: starting conditions 5% ACN, 0.1% formic acid; 1-35 min, 5–50% ACN, 0.1% formic acid; 35-37 min, 50–95% ACN, 0.1% formic acid; 37-40 min 95% ACN, 0.1% formic acid followed by equilibration to starting conditions for additional 20 min. For all runs, we have injected 0.5 μl (0.5μg of human serum proteins derived from 7.1 nl of serum) of tryptic digest directly on chromatography column. We have used a Parallel Reaction Monitoring (PRM) workflow with one MS1 full scan (400-1800 m/z) and scheduled MS/MS fragmentation of IgG1 glycopeptides either completely cleaved or with one missed cleavage. We created a PRM list for non-labelled IgG glycopeptides as well as for the labeled glycopeptides. Fragmentation spectra were recorded in the range 300-2,000 m/z, with an isolation window 1.6 Da for interscan calibration and 10 Da for intrascan calibration. Normalized collision energy was set to 11 for low CE fragmentation and 35 for high CE fragmentation. MS/MS spectra were recorded with a resolution of 30,000 and MS spectra with a resolution of 120,000. We used the same parameters for the methodlogy of EThcD fragmentation where we used calibrated reaction times and supplemental NCE was set to 11. Measurement of 5 replicates was used for fragmentation characteristic determination.

### Optimization of the LC-MS/MS micro-flow measurement

Glycopeptide separation was achieved on an Ultimate 3000 nanochormatograph in microflow mode using a PEPmap300 capillary column 75 μm × 2cm, 5 μm, 300 Å (Thermo) interfaced with an Orbitrap Fusion Lumos (Thermo). Glycopeptides were separated as follows: starting conditions 2% ACN, 0.1% formic acid; 0-1 min 2%ACN, flow 5μl 1-2 min, 2–5% ACN, 0.1% formic acid, flow 1.5μl; 2-5 min, 5–98% ACN, 0.1% formic acid, flow 1.5μl; 7-9 min 98% ACN, 0.1% formic acid, flow 1.5μl followed by equilibration to starting conditions for an additional 4 min. Microflow multinozzle emitter (NEWOMICS) was used as the microflow sprayer. We have used a Parallel Reaction Monitoring (PRM) workflow with one MS1 full scan (400-1800 m/z) and scheduled MS/MS fragmentation of completely cleaved IgG1 glycopeptides as described previously ^23^.

### LC-MS/MS microflow measurement of the samples of Covid 19 infected patients

Serum samples were measured using the optimized microflow method described above. We injected 0.2 μg of the serum protein digest directly on the column. All measurements were done in triplicate.

### Data analysis

Xcalibur and Quan Browser (Thermo) software was used for quantitative data processing. Processing methods were created for ion extraction from each PRM transition in line with our previous observations ^23^. Briefly, PRM transitions of soft fragments (arm loss) were extracted with 20ppm accuracy. Data were processed with no smoothing and chromatogram was visualized using 10 minutes retention time window of expected retention time. Area of integrated peak was used for further data processing. Further data processing and graphing was carried out in Microsoft Excel.

## RESULTS AND DISCUSSION

### Synthesis of the isotope-labeled IgG glycopeptide standards

In this study, we report a highly convergent and streamlined chemoenzymatic approach for the synthesis ^13^C-labeled fucosylated and non-fucosylated glycopeptides standards for quantitation of IgG glycoforms. The key procedure of this modular approach was the efficient synthesis of ^13^C-labeled IgG1-Fc peptide-GlcNAcFuc glycopeptide by a fucoligase AlfCE274A ^36, 37^, which serves as the key acceptor to afford all fucosylated glycopeptides. It overcomes the substrate specificity limitation of the α1,6-fucosyaltransferase (FUT8), which strictly requires the presence of GlcNAc at α1,3 arm of N-glycan substrate for fucose transfer^38,39^. Afterwards, respective N-glycan could be transferred to the precursors from a corresponding N-glycan oxazoline by a glycosynthase-catalyzed reaction to afford the ^13^C-labeled Fc glycopeptide. Non-fucosylated glycopeptides were transferred with EndoCC-D180H mutant ^40^ while core fucosylated glycopeptide was synthesized with EndoF3-D165A mutant ^41^.

The synthetic route of ^13^C-labeled IgG1-Fc-GlcNAc(Fuc) glycopeptides is depicted in **Scheme 1.** Among possible sites for isotope labeling, we chose ^13^C-labeled GlcNAc (**1**, Cambridge Isotope Laboratories, Inc) as the starting material to incorporate ^13^C-labeling in the core GlcNAc-Asn structure which is shared by all Fc N-glycans. The incorporation of the building block in glycopeptides gives a 6 Dalton difference between the “heavy” and “light” isotopic glycopeptides. The synthesis of ^13^C-labeled glycopeptide started with the conversion of ^13^C-labeled GlcNAc (**1**) to the β-glycosyl azide (**2**) via the α-glycosyl chloride and SN2 azide substitution under phase transfer catalysis. Although the large ^13^C-^1^H and ^13^C-^13^C coupling caused the splitting of proton and carbon signals makes the characterization of the product more complicated, a fully assignment of the proton and carbon signals is achieved by COSY and HSQC NMR (see supporting information). Reducing the azido group in **2** by palladium-catalyzed hydrogenation to generate β-glycosyl amine, followed by amide formation with Fmoc-Asn-OtBu using HATU/DIPEA as coupling reagent to give the protected building block **3**, which was then deprotected using formic acid to give the ^13^C labeled building block **4**. This building block was incorporated in the solid-phase peptide synthesis (SPPS) using the Fmoc chemistry on a Rink Amide AM resin (**Scheme 1A**) following our previously reported protocol ^37^ to provide the ^13^C-Fc-peptide-GlcNAc precursor (**5**). The ^13^C-labeled peptide-GlcNAcFuc precursor (**7**), was readily synthesized by using the fucoligase AlfC-E274A^36^, with fucosyl fluoride (**6**) as the donor (**Scheme 1A**).

### Preparation of different glycans from an egg yolk sialylglycopeptide

We prepared different biantennary complex type N-glycan from a sialylglycopeptide (SGP, **8**) purified from egg yolk powder as shown in **Scheme 1B**. After partial desialylation with 0.5% of trifluoroacetic acid (TFA), SGP was cleaved with Endo-S2 endoglycosidase ^39^, and then separated by anion exchange according to the sialylation status, resulting in S2G2 (**9**), S1G2 (**10**), and G2 (**11**) glycans. S1G2 glycan was processed with a β1,4-galactosidase to get S1G1 glycan (**13**), followed by full desialylation with 0.5% TFA to generate the G1 glycoform (**14**). In parallel, mixture of the G2 glycan and peptide-GlcNAc (**12**) was separated by cation exchange, in which the pep-Gn with two positively charged lysine was captured by SP column. Purified G2 was trimmed to afford a G0 glycan (**15**) with a β1,4-galactosidase. With this process, we could easily prepare 30 to 50 mg S2G2, S1G2, G2, G1, and G0 (most abundant form of IgG) from 500 mg of the SGP. The glycans were converted to oxazolines (**16-20**) for the subsequent chemoenzymatic transfer to the peptides (**Scheme 1B**).

**Scheme 1.**
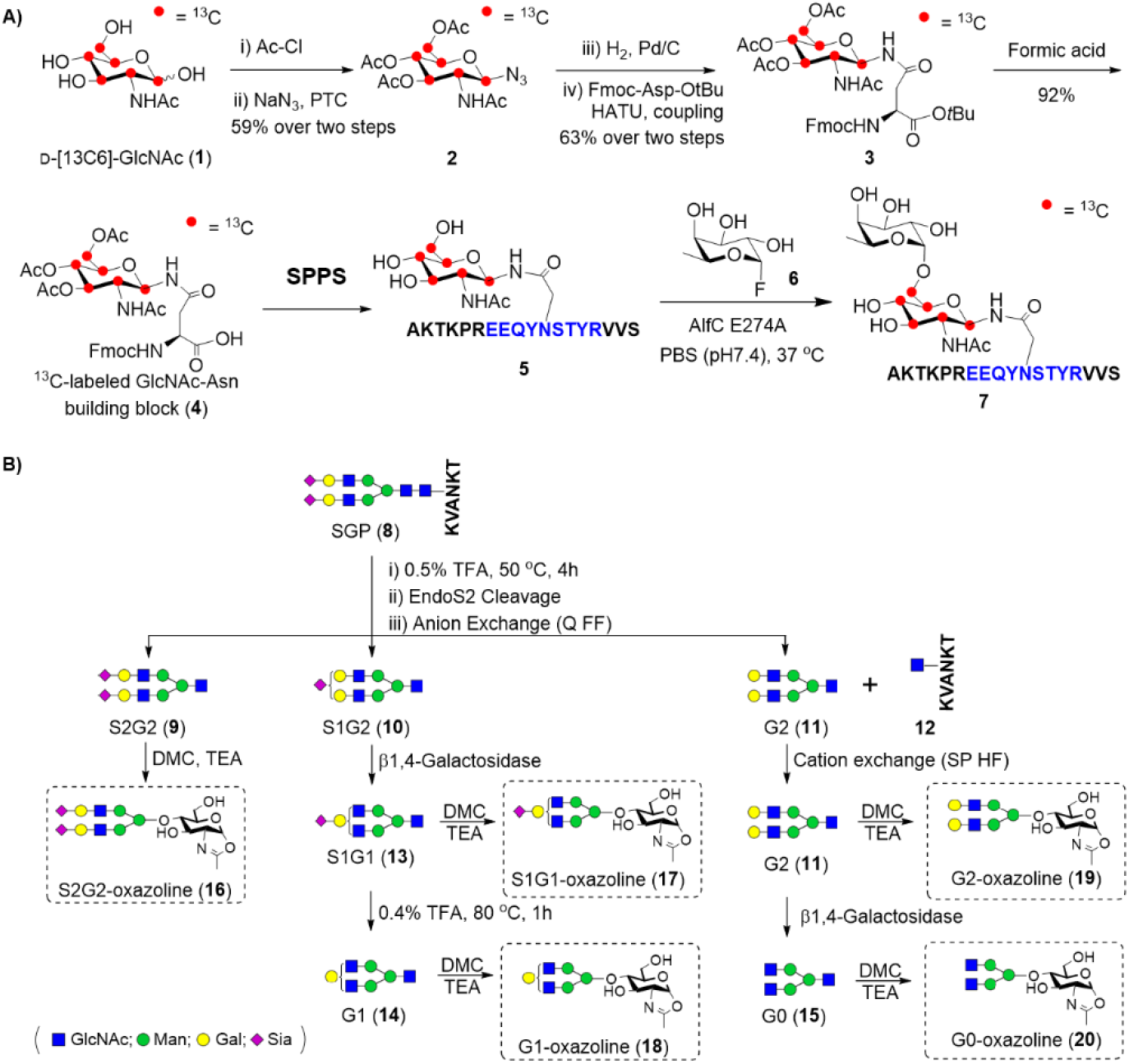
Synthesis of the ^13^C-labeled N-acetylglucosamine (GlcNAc)-peptide precursors (A) and different N-glycan oxazolines (B).

### Synthesis of C-labeled IgG1 Fc glycopeptide library

A ^13^C-labeled IgG1 Fc glycopeptide library was prepared by glycosynthase-catalyzed transfer of a corresponding N-glycan oxazoline to the peptide precursors (**Scheme 2**). Non-fucosylated glycopeptides (**21, 23–26**) were transferred with EndoCC-D180H mutant which is more stable than the EndoM-N175Q mutant (another glycosynthase that transglycosylates complex glycans to the non-fucosylated acceptors) and generally give >95% transfer efficiency. Core fucosylated glycopeptides (**22, 27–30**) were synthesized with EndoF3-D165A mutant which also achieves >95% glycan transfer in all cases tested. The glycopeptide product was purified with semi-preparative HPLC and characterized with LC-ESI-MS. **Table 1** shows a summary of the IgG1 Fc peptides (Fc1P) synthesized. The HPLC and ESI-MS profile of each glycopeptide is shown in **Supplementary Figure 4 and 5**. The ^13^C-labeled peptide was quantitated by UV absorbance at 280 nm, using a non-isotope labeled Fc1P-Gn peptide as the internal standard (**Supplementary Figure 6**).

**Scheme 2.**
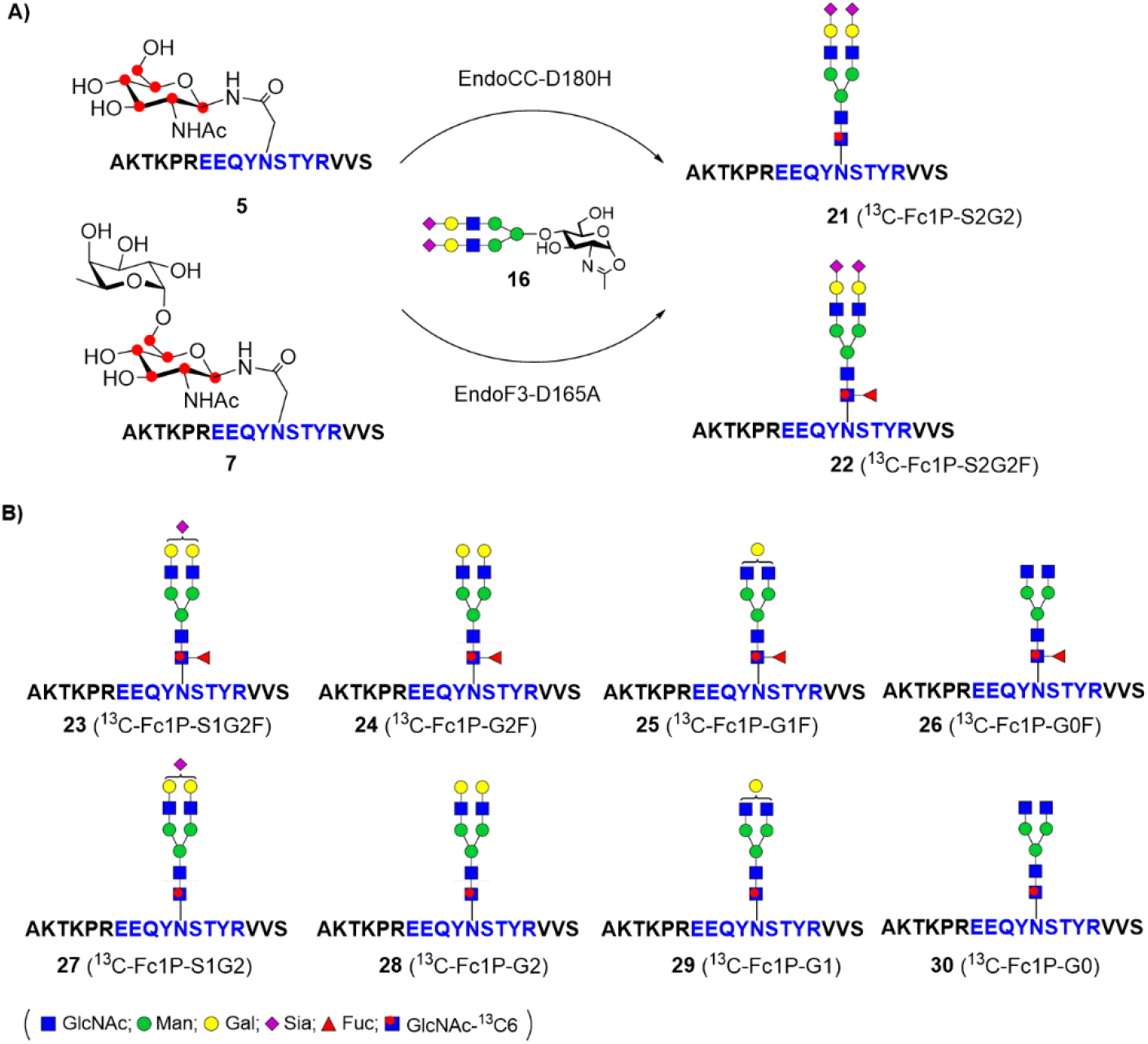
Chemoenzymatic synthesis of various isotope labelled IgG1-Fc glycopeptides.

**Table 1.** Synthesized ^13^C-labeled IgG1 glycopeptides.

| Peptide | M.W. | Quantity (mg) | Yield | Purity (HPLC) |
| --- | --- | --- | --- | --- |
| $^{13}\text{C}$-Fc1P-S2G2 (21) | 4366.28 | 1.49 | >90% | >90% |
| $^{13}\text{C}$-Fc1P-S1G2 (27) | 4075.19 | 1.64 | >90% | >90% |
| $^{13}\text{C}$-Fc1P-G2 (28) | 3784.09 | 0.73 | >90% | >95% |
| $^{13}\text{C}$-Fc1P-G1 (29) | 3622.06 | 1.31 | >90% | >95% |
| $^{13}\text{C}$-Fc1P-G0 (30) | 3459.99 | 1.14 | >90% | >95% |
| <sup>13</sup>C-Fc1P-G0F (26) | 3606.04 | 1.55 | >90% | >95% |
| <sup>13</sup>C-Fc1P-S2G2F (22) | 4512.33 | 1.08 | >90% | >95% |
| <sup>13</sup>C-Fc1P-S1G2F (23) | 4221.24 | 1.75 | >90% | >95% |
| <sup>13</sup>C-Fc1P-G2F (24) | 3930.17 | 1.72 | >90% | >95% |
| <sup>13</sup>C-Fc1P-G1F (25) | 3768.09 | 1.49 | >90% | >95% |

### Fragmentation of IgG standards using low and high collision energy (HCD) fragmentation

We optimized the fragmentation of a core fucosylated synthetic glycopeptide under several acquisition conditions. Under low collision energy CID on a Sciex Q-TOF 5600 ^20^ and HCD on an Orbitrap Fusion Lumos, we observed two major fragments related to the loss of a singly charged N-glycan arm, with and without mannose, as described previously^20, 35^. Fragmentation of the IgG glycopeptides using CID and HCD resulted in a similar fragmentation profile (data not shown). For final testing of selectivity and reproducibility we used HCD with low (11) and high (35) NCE as well as narrow (1.6 Da) and wide (10 Da) window. Fragmentation was tested on purified IgG glycopeptide standards, while selectivity of the signals for IgG MS/MS product ions was studied using serum spiked with a mixture of the IgG glycopeptide standards.

### Quantification of the IgG glycopeptides under low vs high NCE conditions

We have compared HCD fragmentation of IgG glycopeptides for PRM quantification in unfractionated human serum using high and a low CE conditions as described previously 20. Briefly, glycopeptides were fragmented to high-intensity B ions (oxonium ions) and medium-intensity Y ions containing peptide-HexNAc and peptide HexNAc-Fuc fragments. Selectivity of the major oxonium ion 366.1 (HexNAc-Hex) and major Y ion (Peptide-HexNAc) for PRM glycopeptide quantification in unfractionated human serum is shown in **Figure 1**. Specificity of the HexNAc-Hex disaccharide produced by the fragmentation of all galactosylated peptides is not sufficient for a selective quantification of the IgG1 glycopetides except the G1 glycopeptide. The more specific Y1 fragment (peptide-HexNAc) provides sufficient signal to noise (S/N) ratio for quantification of the IgG1 glycopeptides. In case of the asymmetric structure, such as G1, the S/N of the Y1 fragment is on the border of the LOQ. A combination of low CE fragmentation with soft fragment monitoring provides the highest intensities and S/N which enables the PRM mass spectrometric analysis of low-abundant IgG1 glycoforms that we could not reach in the unfractionated human serum using the high CE methods (**Figure 1**).

**Figure 1.**
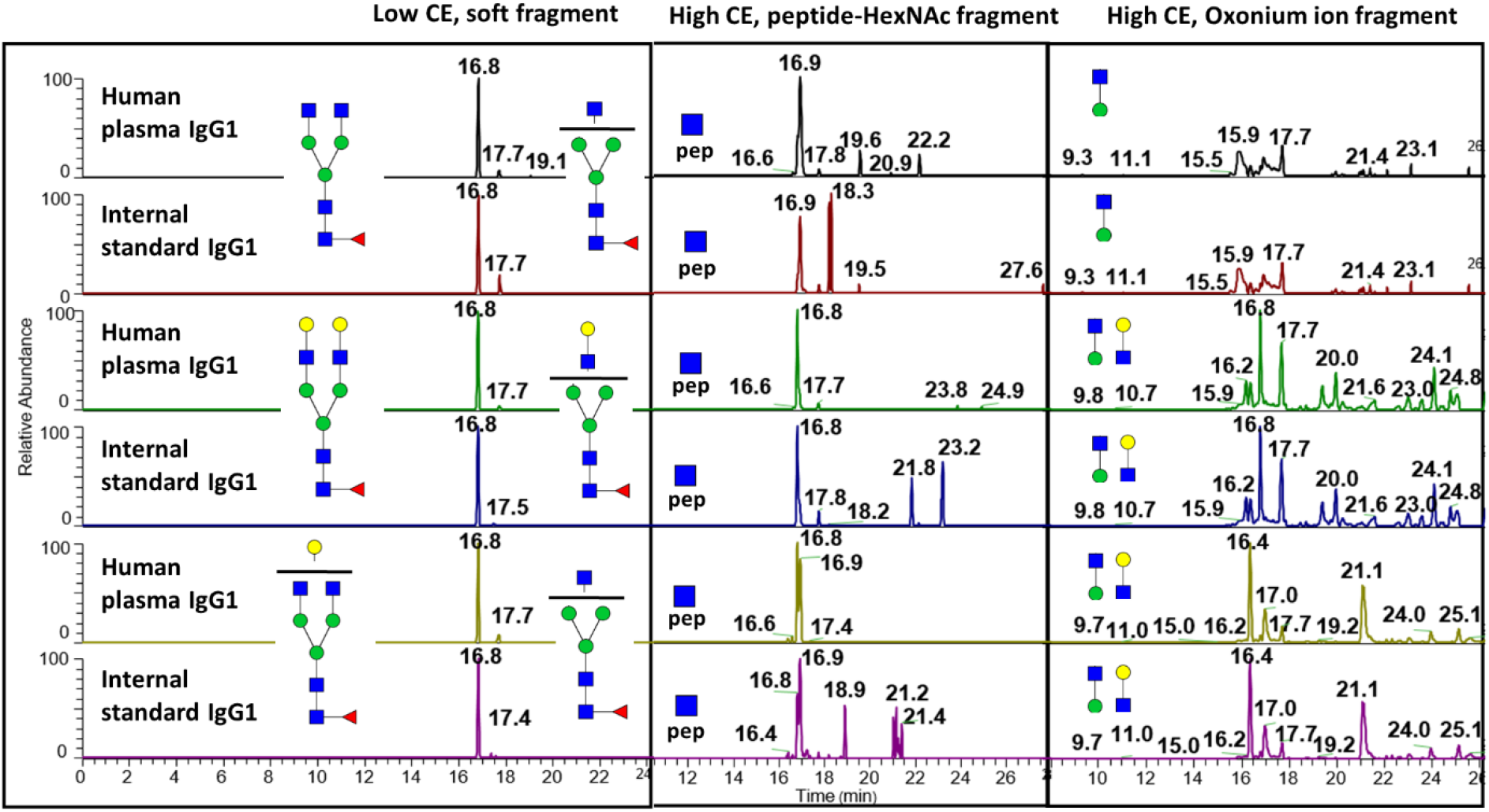
Selectivity of glycopeptide fragments recorded under different CE conditions. Low CE condition (NCE 11), signal of antenna loss Y fragment ion (Left), High collision energy (NCE 35), signal GlcNAc peptide Y fragment ion, High collision energy (NCE 35) signal of HexNAcHex oxonium ion

### Inter-and intra-scan normalization using stable isotope labeled standards to reduce variability of the measurement

The primary purpose of internal calibration is to reduce variability due to the fluctuation of the mass spectrometric signal. We have compared performance of our methodology with and without internal calibration. Using internal calibration, we were able to reduce signal variability below 15% over maximal and for the low CE and below 20% for the high CE methods as determined for five replicates of each measurement in the unfractionated human serum (**Table 2**), in line with the FDA guidelines for LOQ determination in biological mass spectrometry measurements. To improve accuracy of the measurement, we have introduced and tested methods for intra-scan normalization of the PRM workflow. The use of a wide (10Da) window for normalization allowed us to monitor the analyte signal and internal standard in the same fragmentation spectrum. This significantly reduced signal fluctuation due to the fragmentation processes (isolation and fragmentation) as opposed to inter-scan normalization where only pre-fragmentation processes (matrix effect etc.) were normalized. An example of MS/MS product spectra of glycopeptides using a wide fragmentation window was presented in **Figure 2**. We tested narrow and wide fragmentation windows for the high and low CE fragmentation methods. Panel A showed low CE fragmentation spectra of the G0F glycoform of the IgG1 peptide with the loss of one HexNAc as a major soft fragment. Panel B showed a CE spectrum of the G0F glycoform of the IgG1 peptide with the Y1 fragment obtained using a wide isolation window. We used the ratio of the monoisotopic ions of the IgG glycopeptide and the labeled standard for signal normalization. **Table 2** documented a significant reduction of the RSD of the intra-scan normalization; we observe RSDs in the interval 0.6-2.8% under the low CE fragmentation conditions. **Figure 3** showed sensitivity and variability comparison of all optimized workflows for 3 tested glycoforms. The best workflow was found to be the intra-scan normalization using low (11) normalized collision energy which is the workflow with the highest sensitivity for all glycoforms and with the lowest variability in 2 out of 3 glycoforms. Comparison of selectivity was shown in **Figure 4**. Isolation window 10Da recorded under low collision energy had similar performance in analysis of unfractionated human serum as isolation window 1.6Da, which did not introduce any significant interferences that could have negative influence to quantitative performance of optimized methodology.

**Figure 2.**
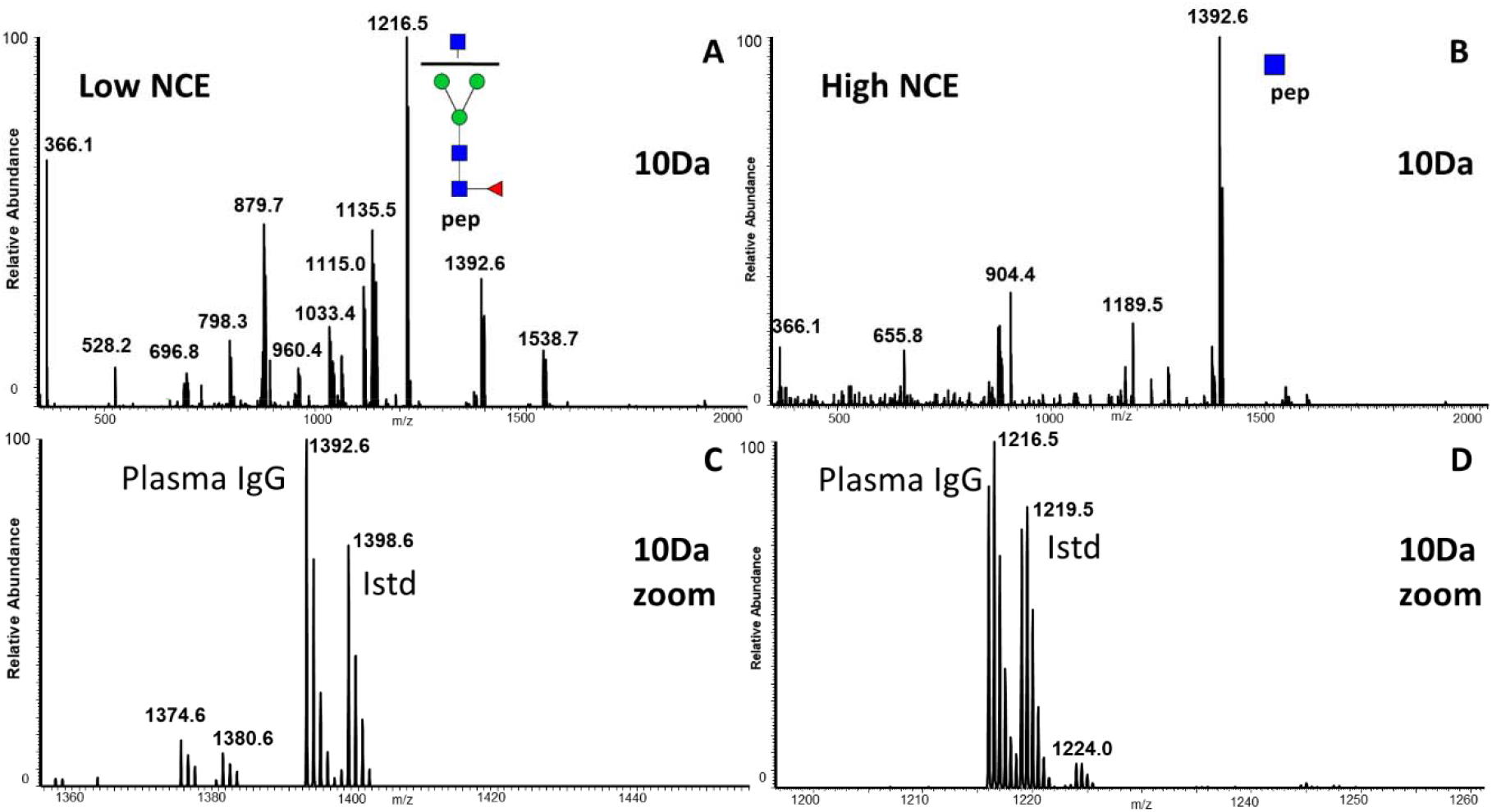
Comparison of the intensities of the most intense soft fragment (low CE) and peptide-HexNAc fragment (high CE) obtained under the following conditions: A. Low NCE, 10 Da window with zoom of qualification ions; B. High NCE, 10 Da window with zoom of quantification ions

**Table 2.** Comparison of the RSD of measurements based on inter-scan (1.6 Da) and intra-scan (10 Da) normalization using unfractionated human serum with spiked labeled IgG internal standards.

| Structure | Low<br>NCE_10 | Low<br>NCE_1.6 | High<br>NCE_10 | High<br>NCE_1.6 | ETHCD<br>10 | ETHCD<br>1.6 |
| --- | --- | --- | --- | --- | --- | --- |
| **G0F** | 2.84 | 5.97 | 5.11 | 36.76 | 12.05 | 43.39 |
| **G2F** | 0.64 | 11.79 | 2.44 | 8.31 | 2.18 | 22.74 |
| **G1F** | 1.24 | 1.97 | 3.11 | 15.30 | 3.20 | 17.68 |

**Figure 3.**
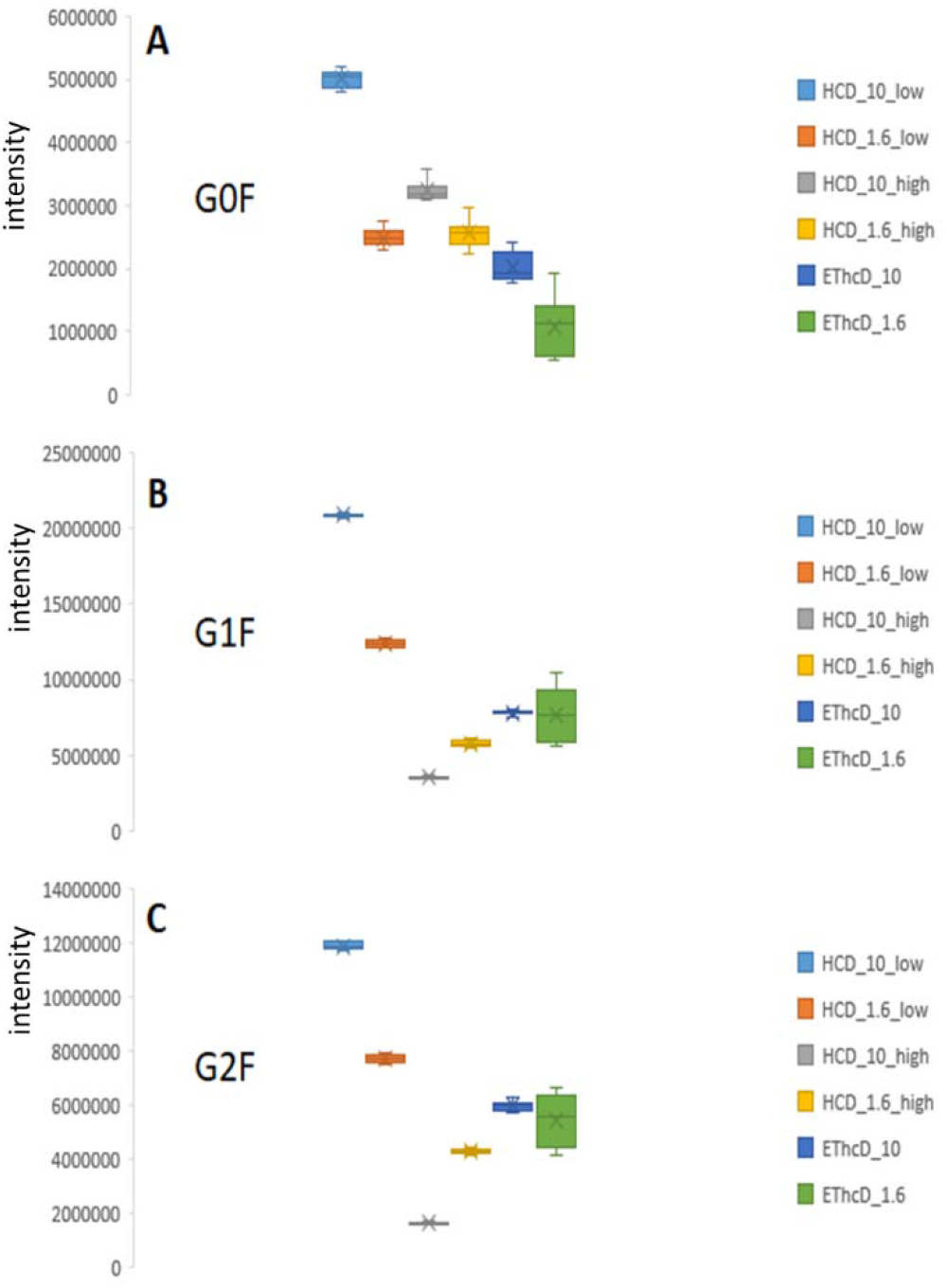
Comparison of the intensities and RSDs of the quantification of three IgG glycoforms (G0F, G1F, and G2F) in the samples of unfractionated human serum. We used HCD with low (11) and high (35) NCE as well as narrow (1.6 Da) and wide (10 Da) window as described in the legend.

**Figure 4.**
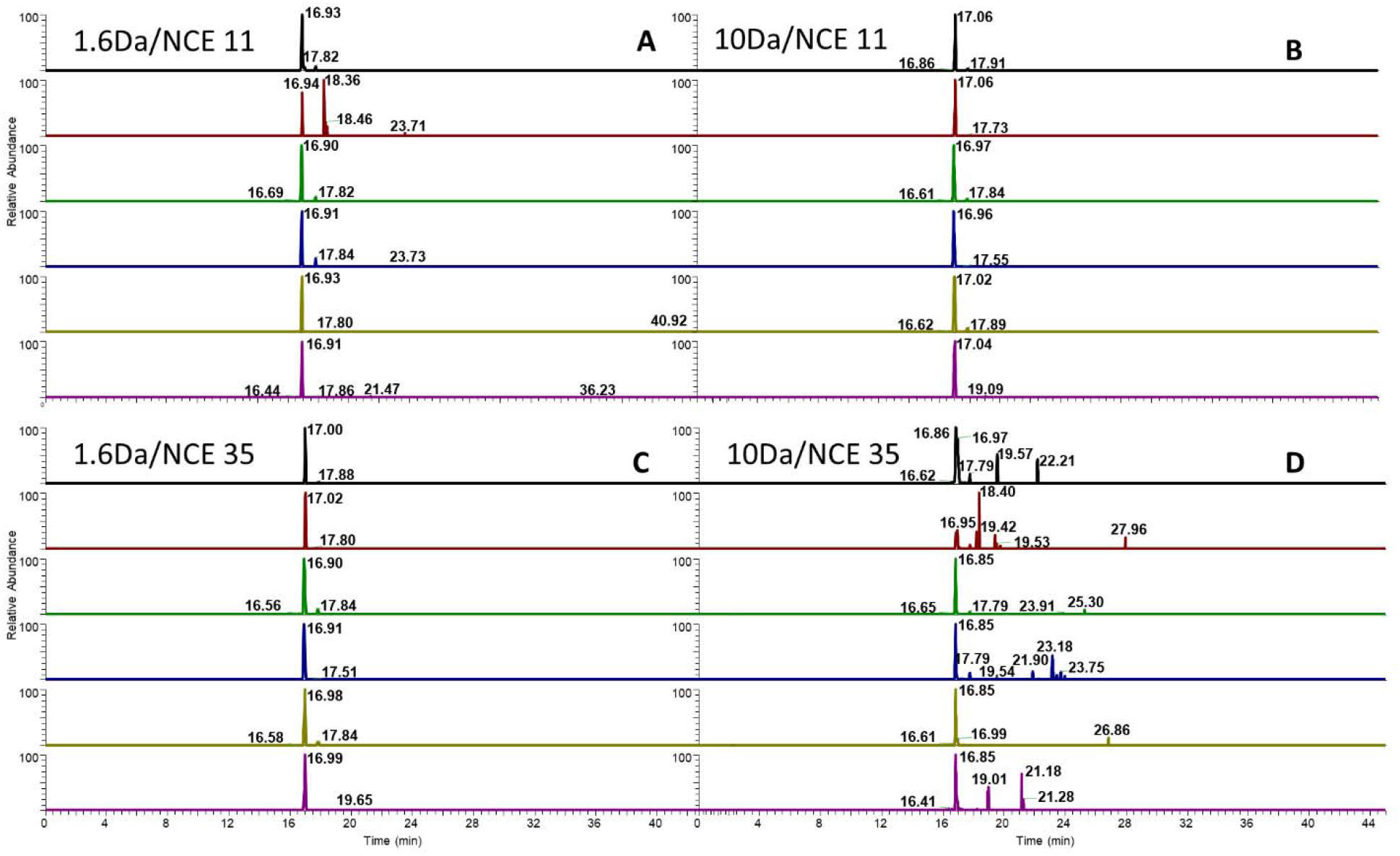
Selectivity of the intra-scan A/C (10Da) and inter-scan B/D (1.6Da) normalization methodology recorded under high XIC signal of peptide-HexNAc Y ion (C/D) and low CE (A/B) signal of antenna loss Y ion.

### A quantitative EThcD workflow

The design of our synthetic IgG standard and high selectivity of the Y-ions allowed us to test a new quantitative application of the EThcD fragmentation workflow even when we use a large isolation window in our PRM workflow. EThcD is primarily used for qualitative analysis like PTM localization on the glycopeptide. As far as we know, a reliable quantitative ETD-based workflow has not been reported yet due to the instability of the inter-scan signal. ETD-based PRM methods could be used to quantify site specific PTMs in case of a mixture of positional isomers. Also, it could be used for quantification of glycopeptides with isobaric peptide backbones like the IgG2 and IgG3 peptides. Therefore, there is a need to develop a robust ETD-based methodology for quantitative mass spectrometric analysis. We used a combination of the wide isolation window with intra-scan normalization to achieve this goal. In this way, we were able to reduce signal variability of the EThcD fragmentation below 15% (**Table 2**) which is in line with the FDA guidelines for LOQ determination in biological mass spectrometric measurements.

### Ultrafast microflow measurement of the IgG glycoforms in a complex matrix

We optimized a fast quantitative PRM workflow which uses microflow chromatography utilizing a multi-nozzle spray. This unique technique enabled analysis of more than 100 samples a day using a 13-minute analytical method. We optimized our method with direct injection onto a 2 cm column and performed a desalting step at a higher flow (5 μl/min) compared to the 2-minute gradient separation at 1.5 μl/min. The equilibration step was performed, again, at a higher flow rate (5 μl/min). A combination of desalting and analytical steps on one column with highly sensitive multi-nozzle spray tip is the key to our fast chromatography with nanoflow like sensitivity. Using our PRM methodology, we were able to get an average of 12 points per chromatographic peak (20s) which exceeds the 8 points per peak recommended for a reliable quantitative analysis. **Supplemental Figure 7** showed an XIC chromatogram of the IgG1 glycoforms analyzed by the optimized microflow method. We also optimized the amount of injected sample with the aim to maximize sensitivity of the method. We determined that maximum sensitivity for quantification of the IgG1 glycoforms could be achieved with an injection of 0.2 μg of an unfractionated serum digest on column. This observed optimum (**Supplemental Figure 8**) results probably from matrix effects related to co-eluted interferences affecting the ionization process.

### IgG1 glycosylation changes in COVID-19 disease

As a practical example of using our optimized methodology, we analyzed IgG1 glycoforms in the serum of healthy volunteers (M, n=5) and COVID 19 patients with severe (S, n=6) conditions (Supplemental Table 1). We quantified 19 previously reported glycoforms of the IgG1 peptide using the microflow LC-MS/MS PRM workflow. We performed the measurement in triplicates. Reproducibility of our measurement using normalized intensities was mostly below 10% (Supplemental Table 2). Despite the precision of our measurement, we did not observe any significant quantitative differences between the M and S groups, either for the 19 individual glycoforms determined or the calculated ratios of glycoforms related to fucosylation, bisecting glycan, sialylation and galactosylation. This observation is in line with previously reported results ^43, 44^. The smaller changes s in the quantified glycoforms of the total pool of antibodies compare to covid specific antibodies likely means that enrichment of the CoV2 specific antibodies is needed to observe the disease-related changes in IgG glycosylation ^43^. In summary, our microflow LC-MS/MS-PRM workflow with the newly available SIS standards achieves sensitive and accurate quantification of the IgG glycoforms in unfractionated serum using a 13-minute workflow. The normalization using the SIS standards reduces the coefficients of variability of the quantification of the glycoforms to less than 5%. We demonstrate that the combination of the wide isolation window with intra-scan normalization allows EThcD-based fragmentation with signal variability less than 15%.

## Acknowledgments

This work was supported in part by the National Institutes of Health (NIH grants U01CA230692, R01CA238455 and S10OD023557 to RG; R43GM128547 to QY; and R01GM080374 to LXW; and R01AI132389 to AK). The content is solely the responsibility of the authors and does not necessarily represent the official views of the National Institutes of Health.

