## Supplemental material for "LC-MS/MS-PRM Quantification of IgG glycoforms using stable isotope labeled IgG1 Fc glycopeptide standard"

^a^Department of Oncology, Lombardi Comprehensive Cancer Center, Georgetown University, Washington, D.C. 20057; ^b^GlycoT Therapeutics, College Park, MD 20742, ^c^Department of Chemistry and Biochemistry, University of Maryland, College Park, MD 20742, Georgetown University, Washington, D.C., 20057; ^d^MedStar Georgetown Transplant Institute, MedStar Georgetown University Hospital and the Center for Translational Transplant Medicine, Georgetown University Medical Center, Washington, D.C. , Washington, D.C., 20057; ^e^Clinical and Translational Glycoscience Research Center, Georgetown University, Washington, D.C., 20057; ^f^Department of Biochemistry and Molecular & Cell Biology, Georgetown University, Washington, D.C., 20057, Max-Planck-Institut fuer Herz- und Lungenforschung, Ludwigstrasse 43, Bad Nauheim, 61231, Germany ; †Authors contributed equally to this work.

Table of content

1. NMR spectrum of compound **2-4**
2. RP-HPLC and ESI-MS analysis of synthesized ^13^C-labeled IgG1-Fc-GlcNAc
3. RP-HPLC (A) and ESI-MS (B) analysis of synthesized ^13^C-labeled fucosylated IgG1-FcGlcNAcFuc.
4. RP-HPLC and ESI-MS analysis of ^13^C-labeled fucosylated IgG1-Fc glycopeptides.
5. RP-HPLC (A-E) and ESI-MS (A’-E) analysis of ^13^C-labeled non-fucosylated IgG1-Fc glycopeptides.
6. RP-HPLC quantitation of the synthesized ^13^C-labeled glycopeptides with unlabeled IgG1-Fc-GlcNAc peptide as the internal standard reference
7. Comparison of HCD (Orbitrap Fusion Lumos) and CID (q-tof) soft fragmentation spectra of the biantennary galactosylated structure of IgG1
8. Extracted chromatograms of the glycoforms of IgG1 using the optimized microflow methodology
9. Optimization of the amount of material injected on column for maximal sensitivity of the analysis
10. Methodology supplement

**Supplementary Figure 1 (A-H)**

**Supplementary Fig 1A.** 1H NMR spectrum of compound **2**

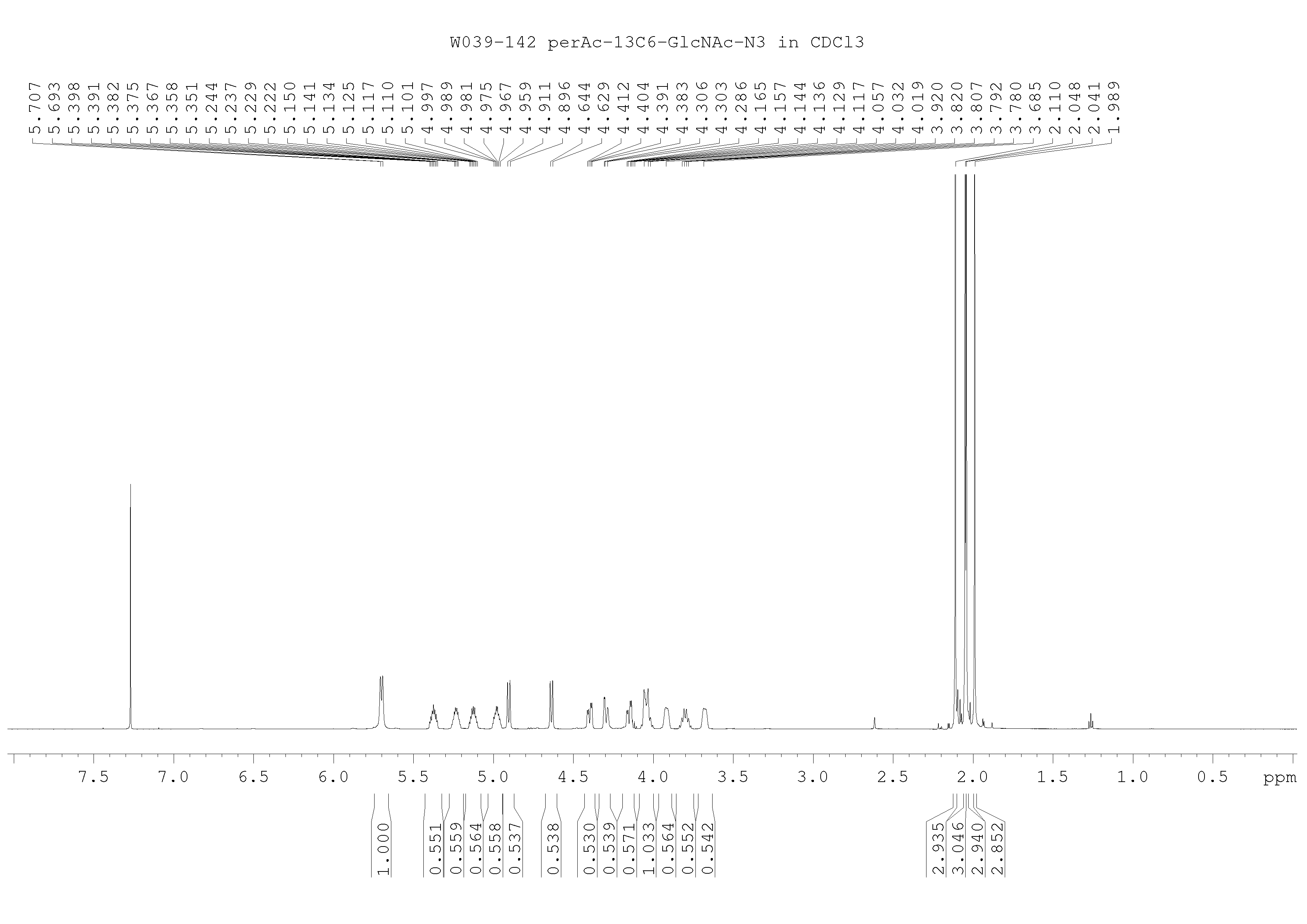

**Supplementary Fig 1B.** 13C NMR spectrum of compound **2**

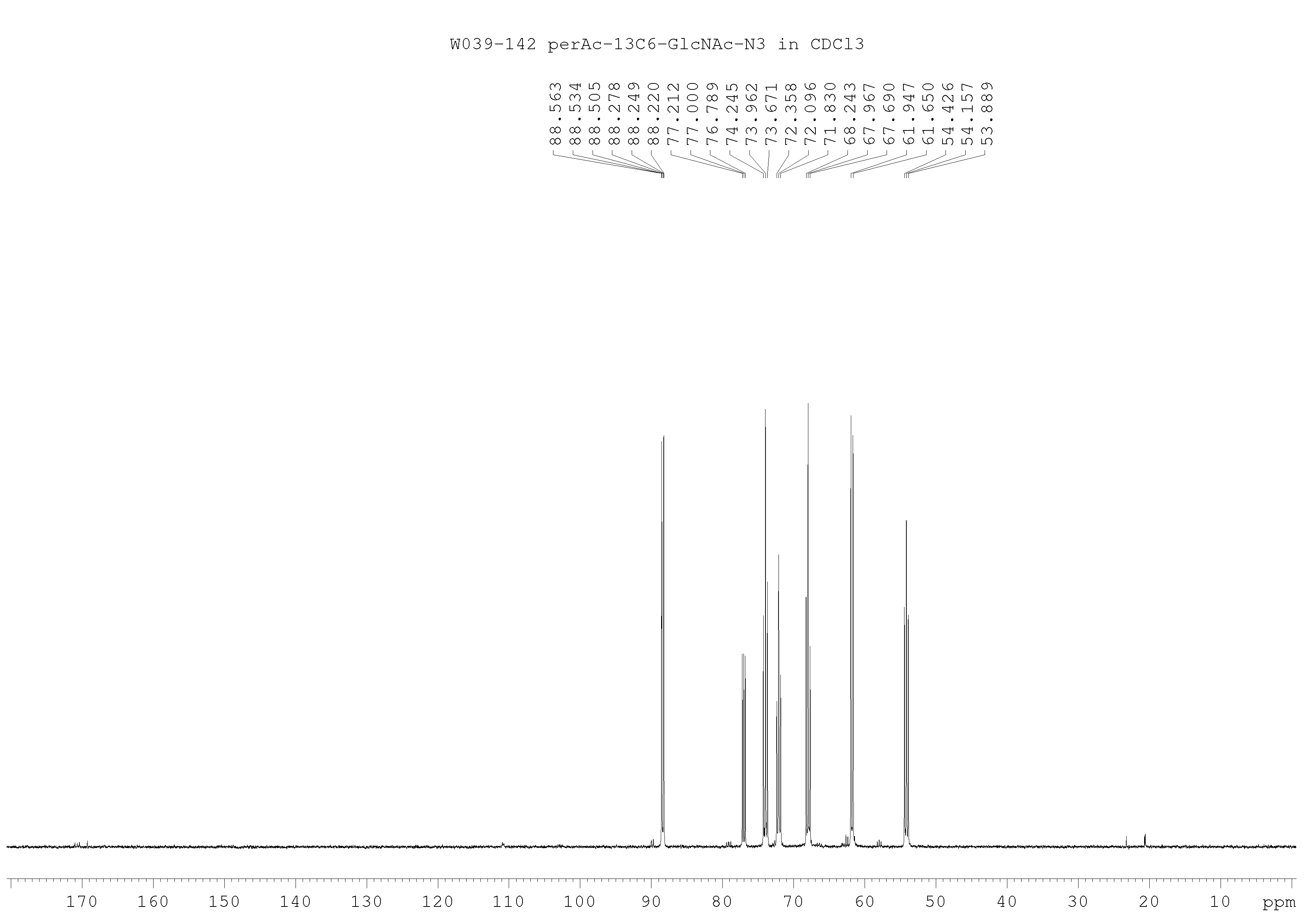

**Supplementary Fig 1C.** COSY NMR spectrum of compound **2**

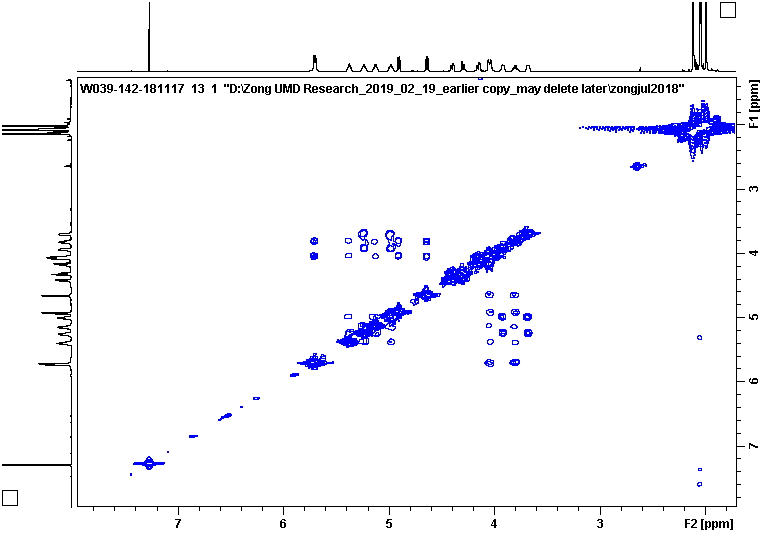

**Supplementary Fig 1D.** HSQC NMR spectrum of compound **2**

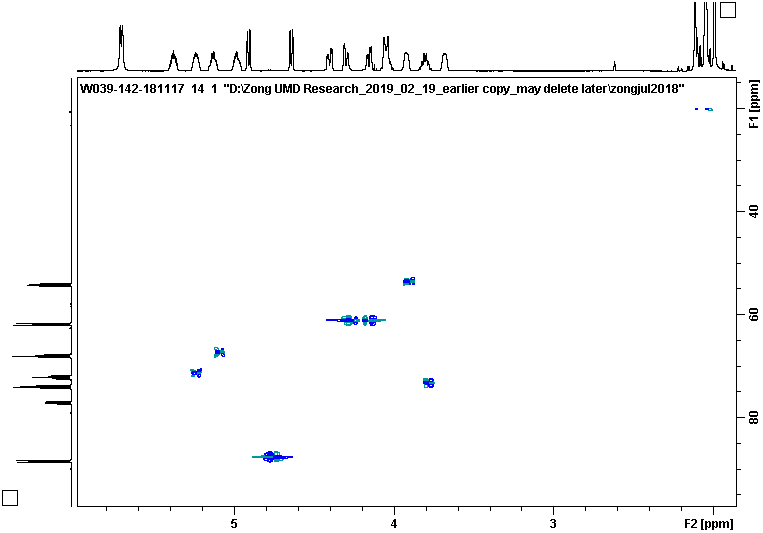

**Supplementary Fig 1E.** 1H NMR spectrum of compound **3**

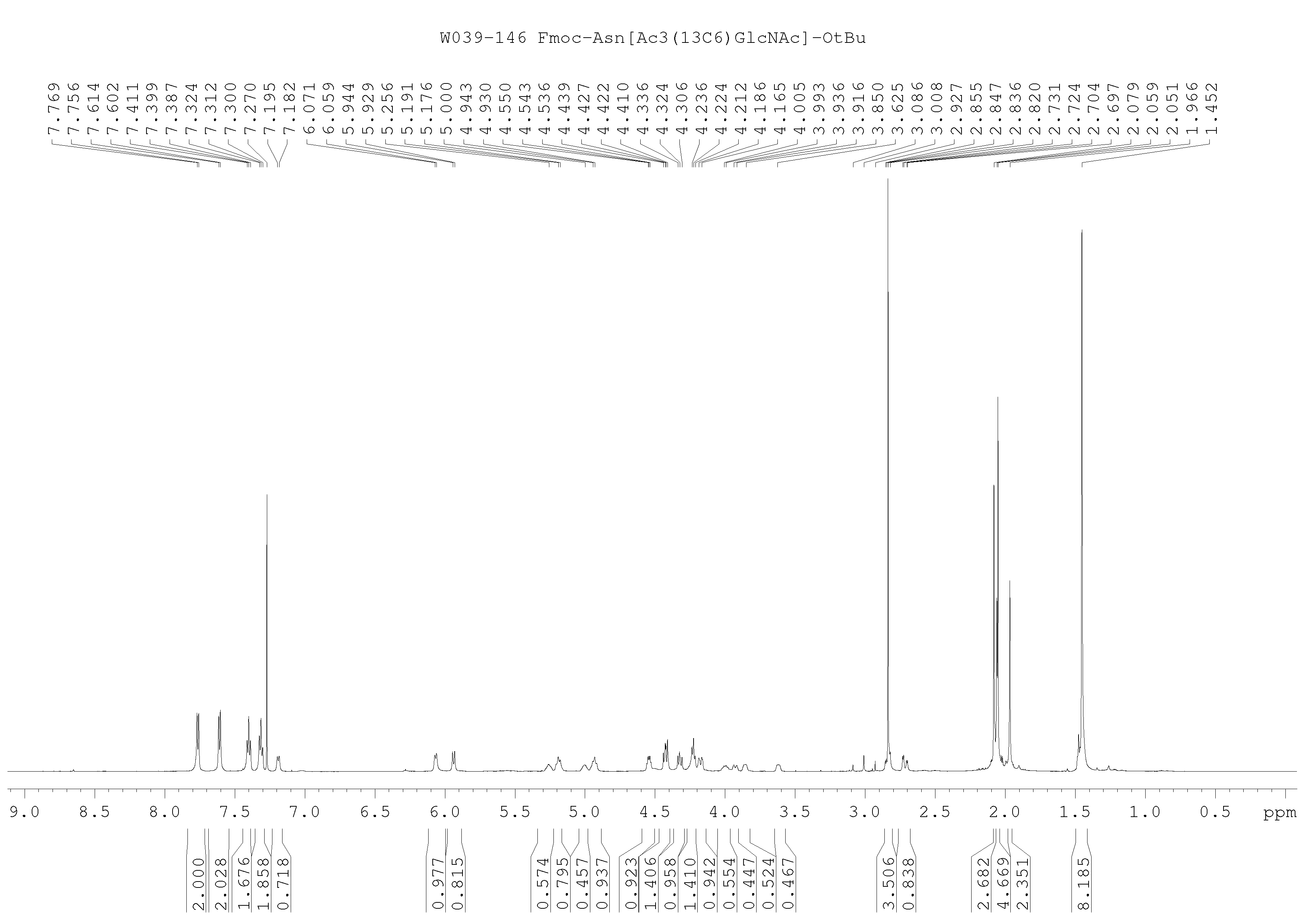

**Supplementary Fig 1F.** 13C NMR spectrum of compound **3**

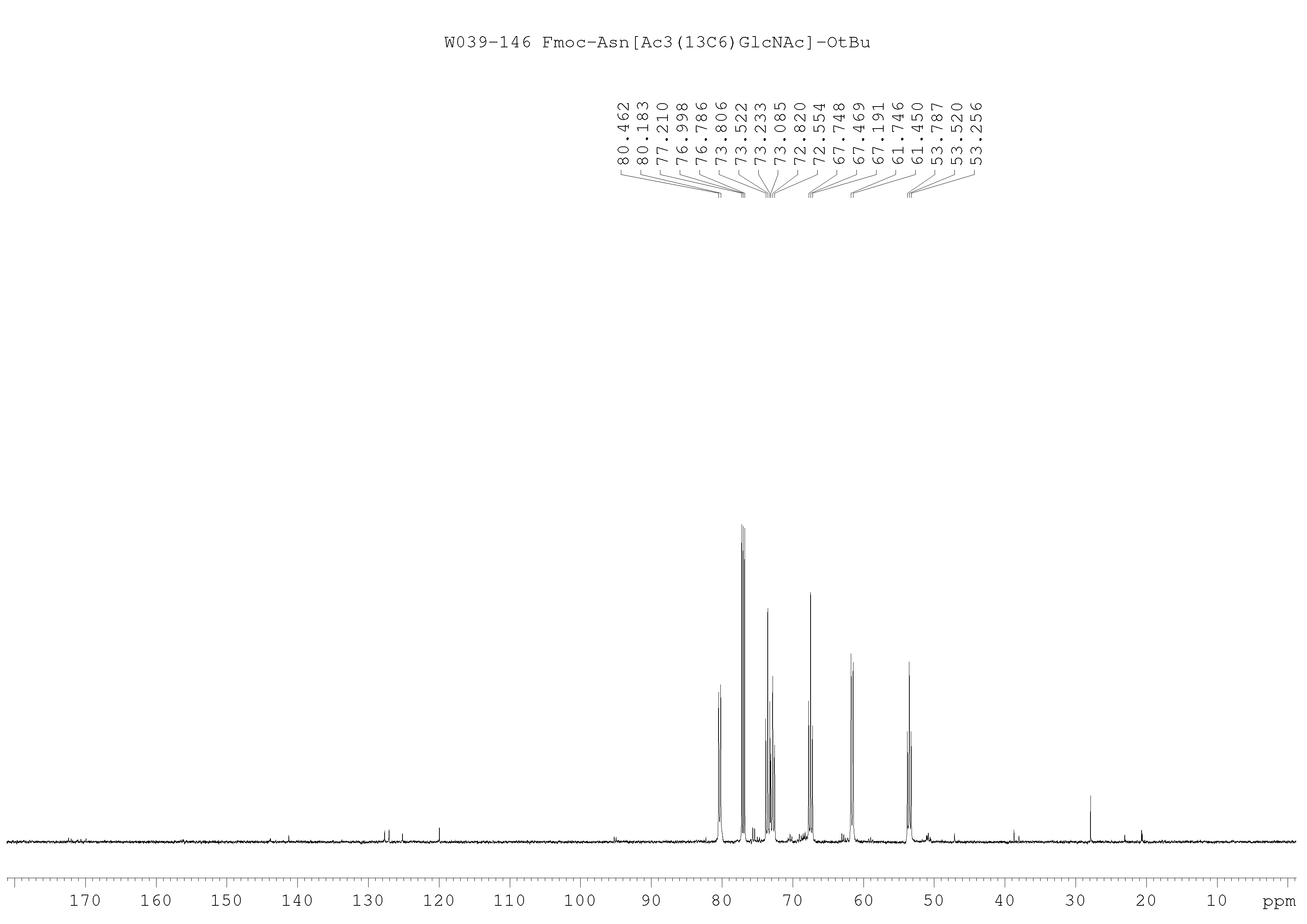

**Supplementary Fig 1G.** 1H NMR spectrum of compound **4**

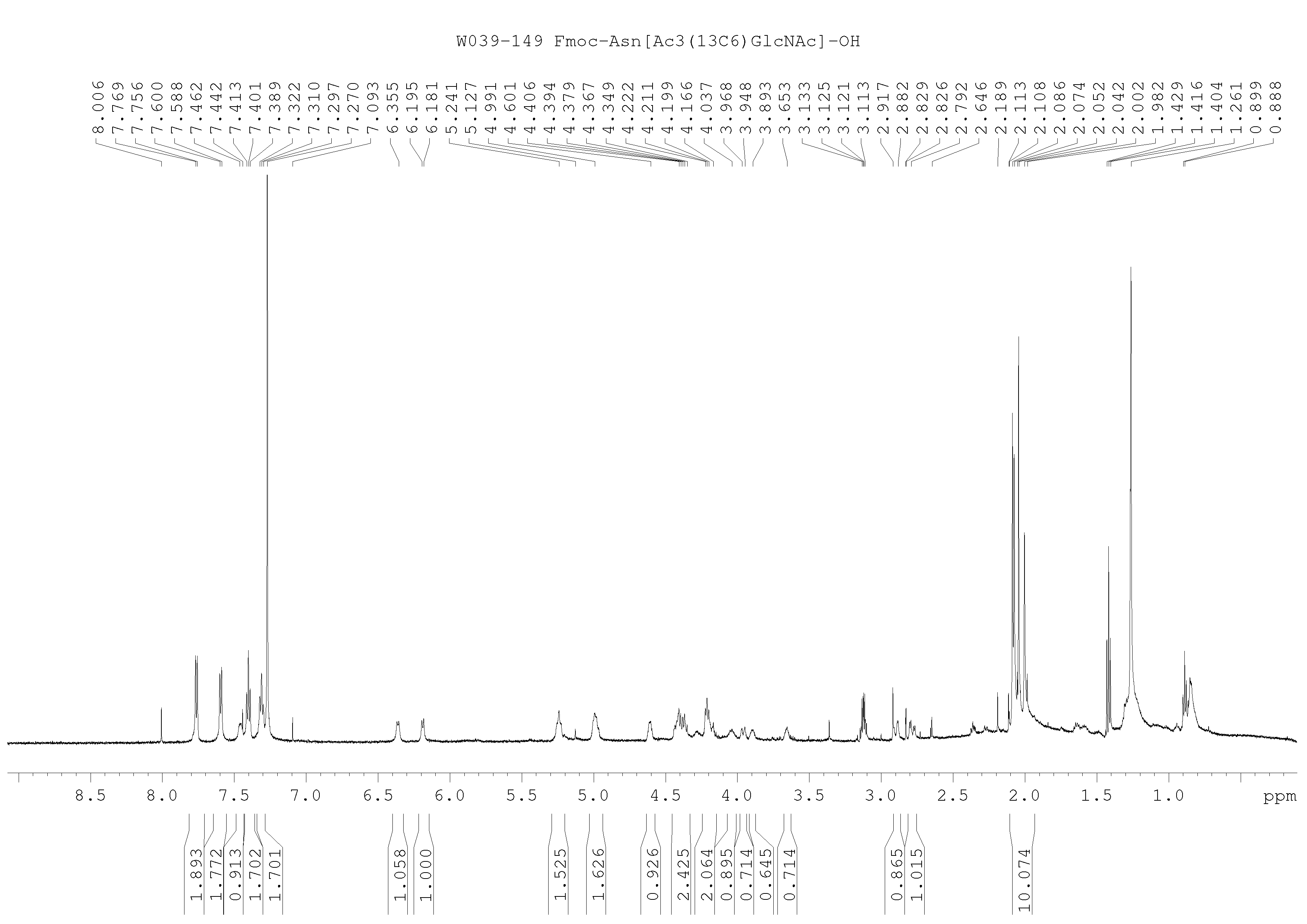

**Supplementary Fig 1H.** ^13^C NMR spectrum of compound **4**

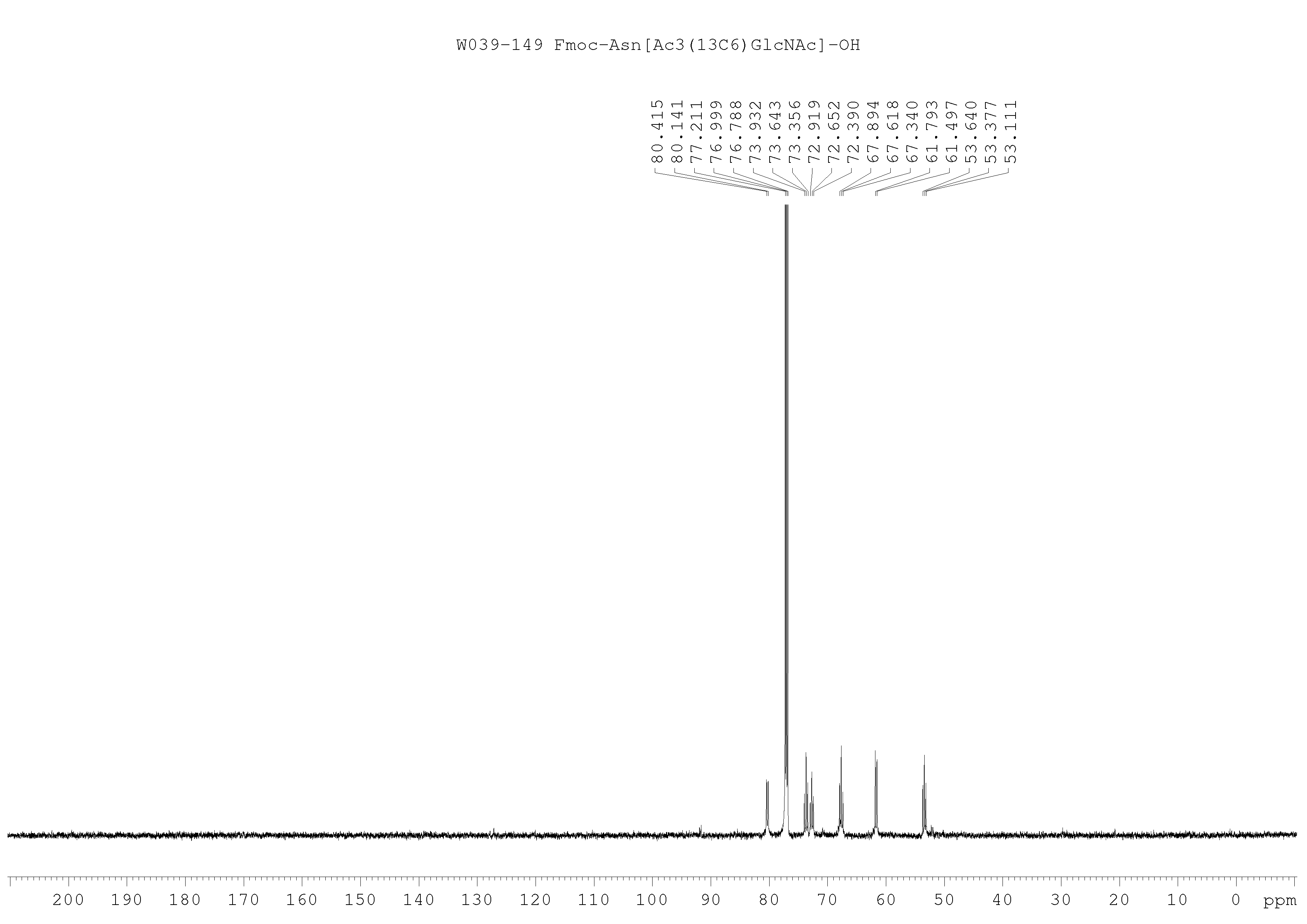

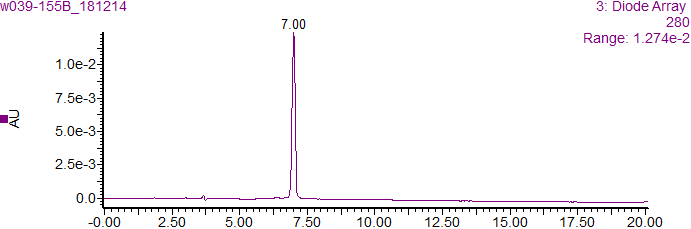

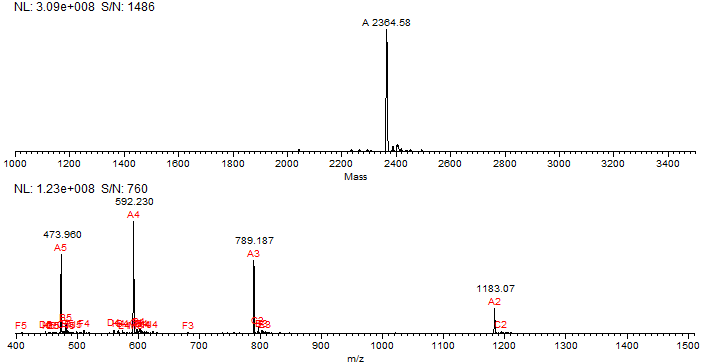

**Supplementary Fig. 2. RP-HPLC and ESI-MS analysis of synthesized ^13^C-labeled IgG1-Fc-GlcNAc (5).**

*GlcNAc-peptide* ***5***. Analytical RP-HPLC, *t*_R_ = 7.0 min (5-35% MeCN containing 0.1% formic acid at a flow rate of 1 mL/min over 20 min, C18 column (YMC-Triart C18, 4.6 × 250 mm, 5 μm)). ESI-MS: calcd., M = 2364.68 (average isotopes); found, 473.96 [M+4H]^4+^, 592.23 [M+4H]^4+^, 789.19 [M+3H]^3+^, 1183.07 [M+2H]^2+^. Deconvolution mass, 2364.58.

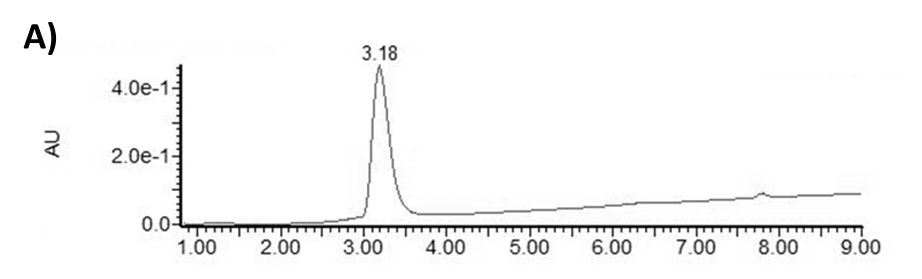

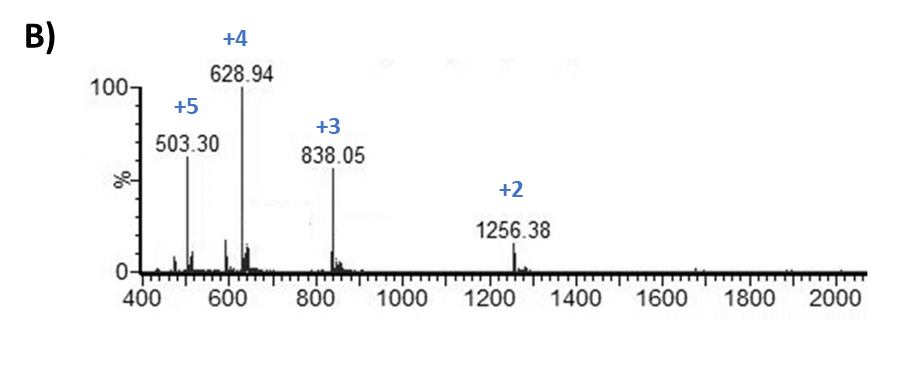

**Supplementary Fig. 3. RP-HPLC (A) and ESI-MS (B) analysis of synthesized ^13^C-labeled fucosylated IgG1-FcGlcNAcFuc (13C-Fc1P-GnF).** Calculated MW: 2510.75

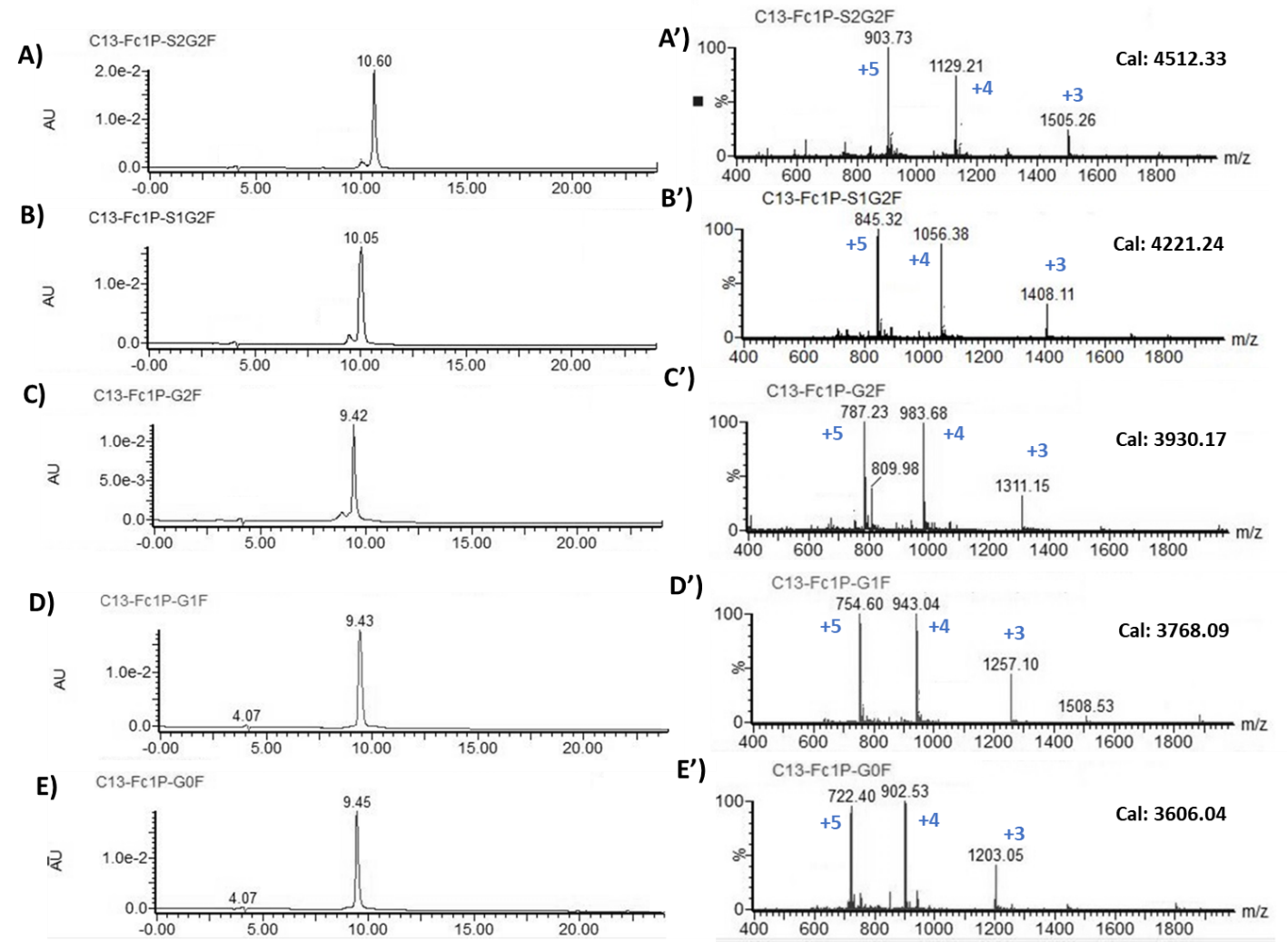

**Supplementary Fig 4. RP-HPLC (A-E) and ESI-MS (A’-E) analysis of ^13^C-labeled fucosylated IgG1-Fc glycopeptides.** Analytical RP-HPLC was performed with a gradient of 5-35% MeCN containing 0.1% formic acid at a flow rate of 1 mL/min over 30 min on a C18 column (YMC-Triart C18, 4.6 × 250 mm, 5 μm).

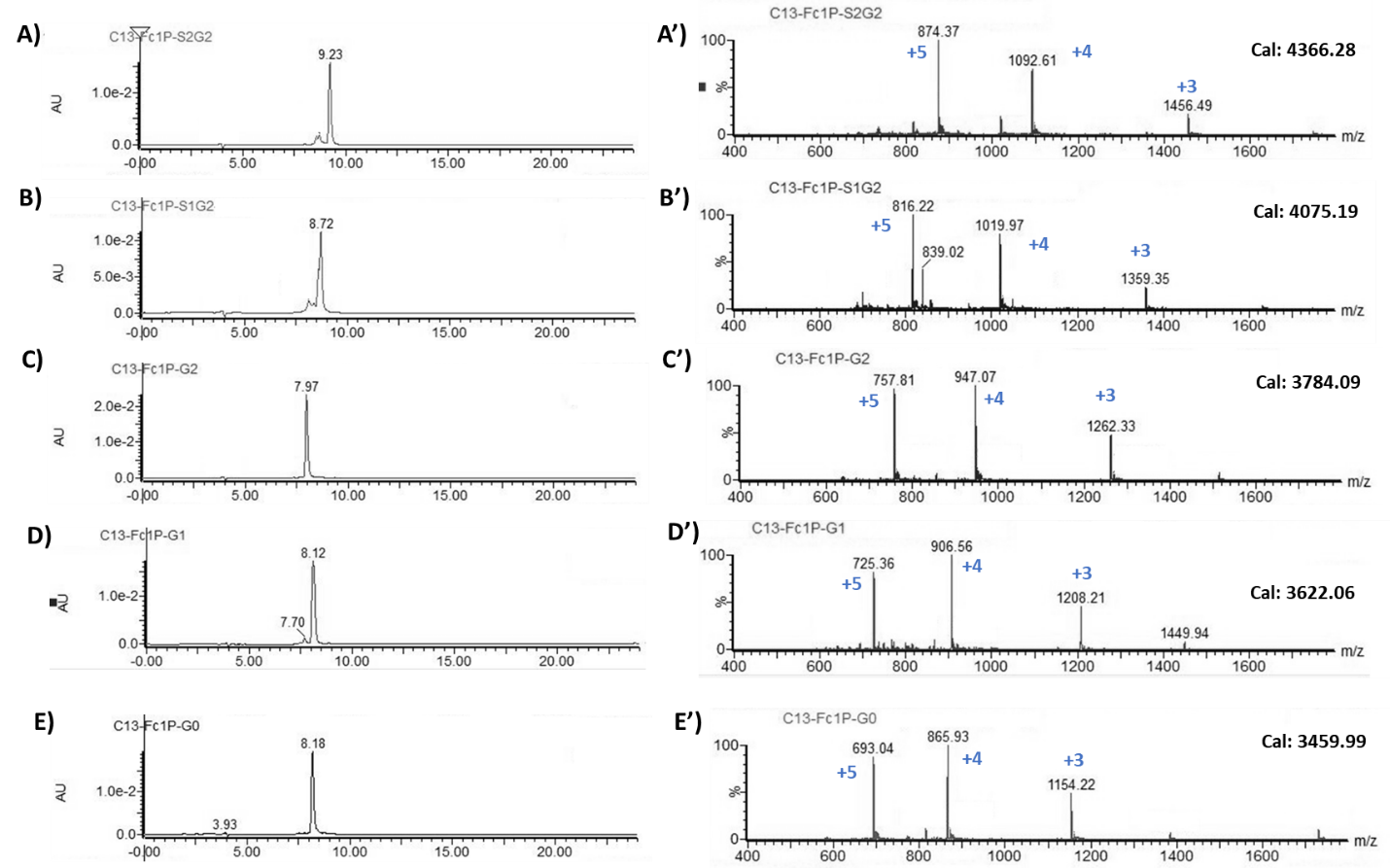

**Supplementary Fig 5. RP-HPLC (A-E) and ESI-MS (A’-E) analysis of ^13^C-labeled non-fucosylated IgG1-Fc glycopeptides.** Analytical RP-HPLC was performed with a gradient of 5-35% MeCN containing 0.1% formic acid at a flow rate of 1 mL/min over 30 min on a C18 column (YMC-Triart C18, 4.6 × 250 mm, 5 μm).

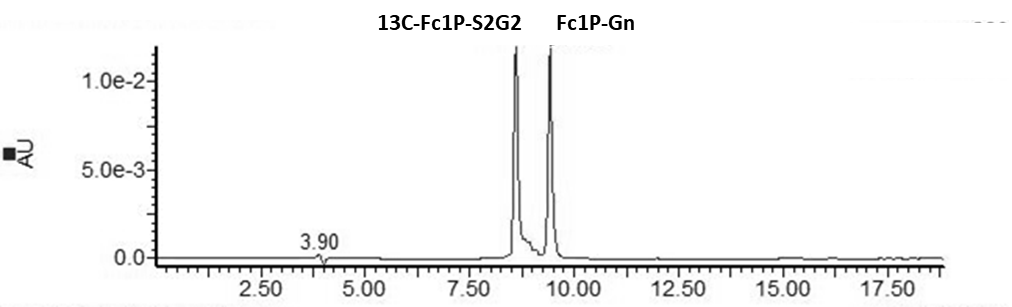

**Supplementary Fig 6. RP-HPLC quantitation of the synthesized ^13^C-labeled glycopeptides with unlabeled IgG1-Fc-GlcNAc peptide (Fc1P-Gn) as the internal standard reference.** HPLC profile of ^13^C-Fc1P-S2G2 with incorporation of Fc1P-Gn internal standard was shown as an example. RP-HPLC was performed with a 5-30% ACN containing 0.1% FA, 20 min gradient, 55 °C on a Xbridge C18 column (4.6x150 mm 3.5 um).

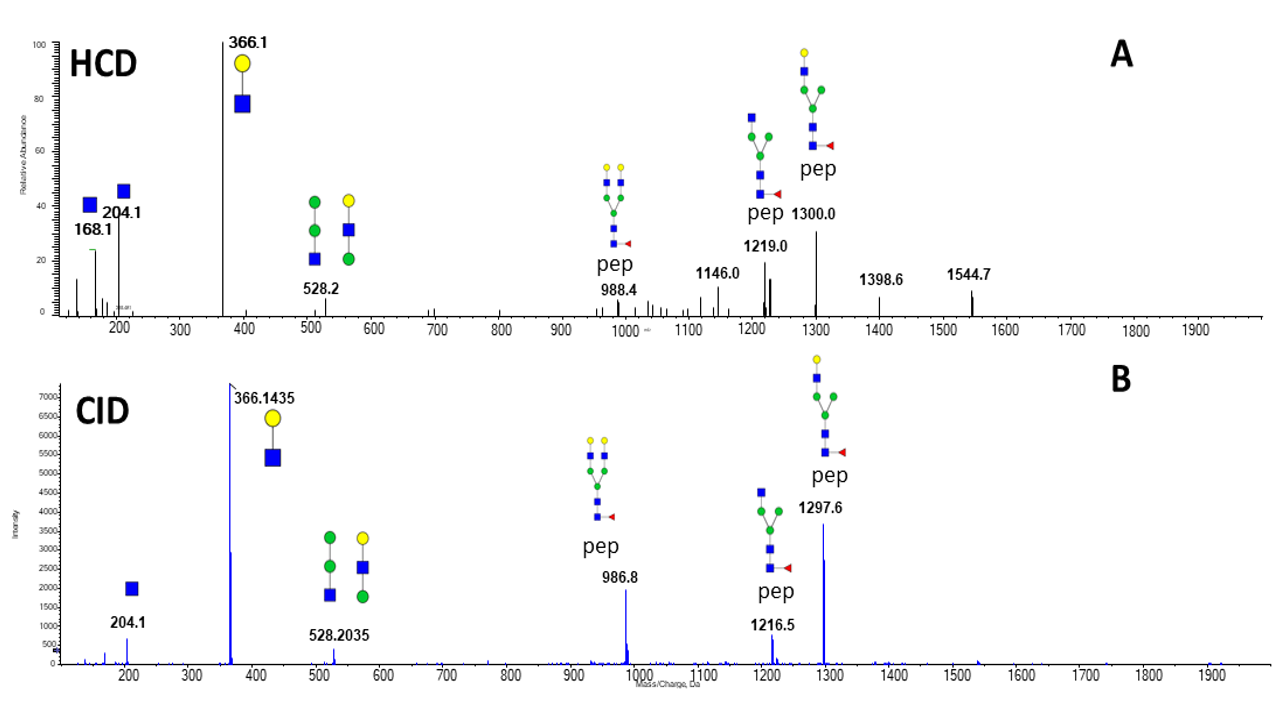

**Supplemental Fig 7.**	Comparison of HCD (Orbitrap Fusion Lumos - stable isotope labeled standard) **A** and CID (q-tof – non-labeled glycopeptide) **B** soft fragmentation spectra of the biantennary galactosylated structure of IgG1

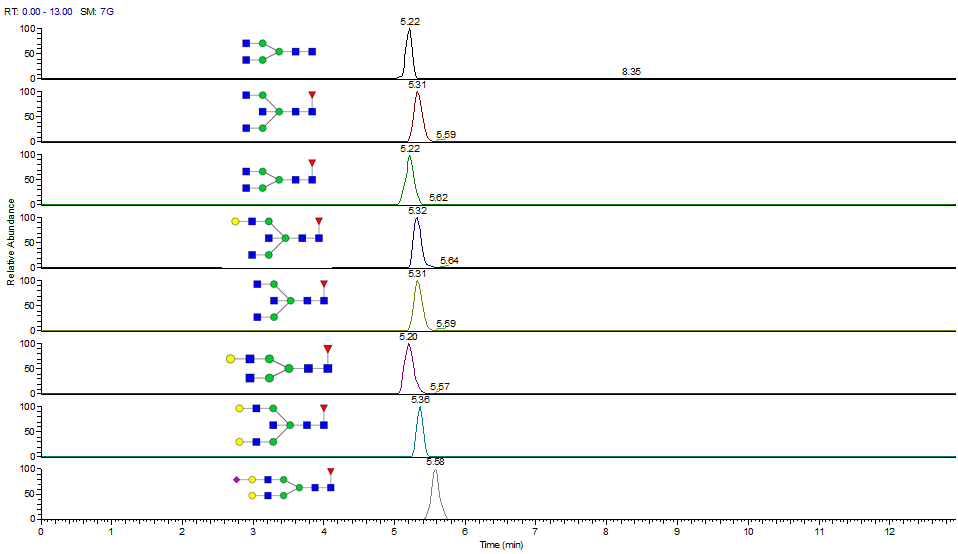

**Supplemental Fig 8.** Extracted chromatograms of the glycoforms of IgG1 using the optimized microflow methodology.

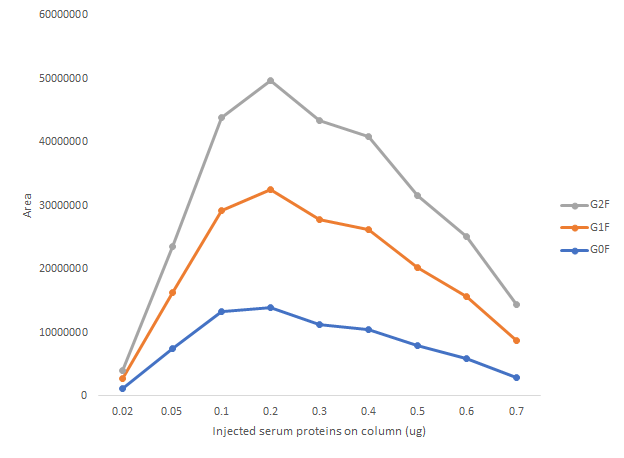

**Supplemental Fig.** **9.** Optimization of the amount of material injected on column for maximal sensitivity of the analysis.

10. Methodology supplement:

**Synthesis of ^13^C-GlcNAc-Asn Building Block**

^13^C Labeled building block Fmoc-Asn(Ac_3_[^13^C6]GlcNAc)-OH was prepared following a similar procedure for the synthesis of the non-isotope labeled building block Fmoc-Asn(Ac_3_GlcNAc)-OH^35^, as depicted in **Scheme 1**.

2-Acetamido-3,4,6-tri-*O*-Acetyl-2-deoxy-*β*-D-[UL-^13^C6]-Glucopyranosyl Azide (2). D-[UL-^13^C6]-N-Acetylglucosamine **1** (600 mg, 2.64 mmol) was added dropwise to cold (0 °C) acetyl chloride (3 mL), and the mixture was stirred atroom temperature for 16 h. The reaction mixture was then diluted with CH_2_Cl_2_ (50 mL) and poured into cold sat. aq NaHCO_3_ with vigorous stirring. The organic layer was separated, dried over anhydrous Na_2_SO_4_, and filtered. The filtrate was concentrated to get crude 2-acetamido-3,4,6-tri-*O*-acetyl-2-deoxy-*α*-D-[UL-^13^C6]-glucopyranosyl chloride as a syrup, which was used for the next step without further purification. To a solution of the above obtained residue, tetrabutylammonium hydrogen sulfate (TBAHS, 0.90 g, 2.64 mmol), and NaN_3_ (0.69 mg, 10.6 mmol) in CH_2_Cl_2_ (20 mL) was added sat. aq. NaHCO_3_ (20 mL). The two-phase mixture was stirred vigorously at room temperature for 2 h, at which time Thin-layer chromatography (TLC) (silica, EtOAc–hexanes, 2:1) indicated complete consumption of the glycosyl halide. The organic phase was separated, and aqueous layer was extracted with CH_2_Cl_2_ (20 mL). The combined organic layer was dried over anhydrous Na_2_SO_4_ and filtered. The filtrate was concentrated and the residue was purified by flash silica gel chromatography (20−100% EtOAc in hexane) to give **2** (590 mg, 59% over two steps) as a white solid. ^1^H NMR (600 MHz, CDCl_3_) δ 5.69 (d, J = 8.4 Hz, NH), 5.44 – 5.32 (m, 0.5H, H-3), 5.28 – 5.19 (m, 0.5H, H-4), 5.17 – 5.08 (m, 0.5H, H-3), 5.03 – 5.94 (m, 0.5H, H-5), 4.77 (dd, J = 9.2, 160 Hz, H-1), 4.44 – 4.36 (m, 0.5H, H-6), 4.34 – 4.27 (m, 0.5H, H-6), 4.18 – 4.12 (m, 0.5H, H-6), 4.08 – 4.00 (m, 1H, H-6, H-2), 3.95 – 3.88 (m, 0.5H, H-5), 3.85 – 3.75 (m, 0.5H, H-2), 3.71 – 3.63 (m, 0.5H, H-5), 2.11 (s, 1H, CH_3_CO), 2.05 (s, 1H, CH_3_CO), 2.04 (s, 1H, CH_3_CO), 1.99 (s, 1H, CH_3_CO). ^13^C NMR (150 MHz, CDCl_3_) δ 88.4 (dt, J = 4.4, 42.8, ^13^C-1), 74.0 (t, J = 43.0, ^13^C-5), 72.1 (t, J = 39.6, ^13^C-3), 68.0 (t, J = 41.4, ^13^C-2), 61.8 (d, J = 44.6, ^13^C-6), 54.2 (t, J = 40.2, ^13^C-2). HRMS (ESI-Orbitrap) m/z [M+H]^+^ Calcd for C^14^H^21^N^4^O^8+^: 379.1555; found, 379.1537.

N-*α*-(Fluorenylmethoxycarbonyl)-N-*γ*-(2-acetamido-3,4,6-tri-*O*-acetyl-2-deoxy-*β*-D-glucopyranosyl)-L-asparagine tert-butyl ester (**3**). To a solution of azide **2** (230 mg, 0.61 mmol) in CH_2_Cl_2_ (10 mL) was added Pd/C (20 mg) in one portion at room temperature. The reaction was then stirred under an atmosphere of hydrogen (1 atm) for 3 h. At this point, TLC (silica, 10:1 CH_2_Cl_2_–MeOH) showed the reaction was complete. The reaction mixture was filtered through a pad of Celite using CH_2_Cl_2_ (10 mL) as the eluent. To the resulting filtrate was added Fmoc-Asp(OtBu)-OH (275 mg, 0.67 mmol), HATU (693 mg, 1.83 mmol) and DIEA (0.32 mL, 1.8 mmol) and the reaction mixture was stirred at rt overnight. Then the reaction mixture was concentrated and the residue was purified by flash silica gel chromatography (0−10% MeOH in CH_2_Cl_2_) to give **3** (285 mg, 63% over two steps) as a white solid. ^1^H NMR (600 MHz, CDCl_3_) δ 7.77 (d, J = 7.4 Hz, 2H, Fmoc-Ar), 7.60 (d, J = 7.4 Hz, 2H, Fmoc-Ar), 7.40 (t, J = 7.4 Hz, 2H, Fmoc-Ar), 7.31 (t, J = 7.4 Hz, 2H, Fmoc-Ar), 7.19 (d, J = 7.6 Hz, 1H), 6.06 (d, J = 7.4 Hz, 1H), 5.93 (d, J = 8.8 Hz, 1H), 5.30 – 5.22 (m, 0.5H), 5.21 – 5.15 (m, 1H), 5.05 – 4.99 (m, 0.5H), 4.98 – 4.89 (m, 1H), 4.58 – 4.48 (m, 1H), 4.47 – 4.38 (m, 1.5H), 4.37 – 4.29 (m, 1H), 4.27 – 4.20 (m, 1.5H), 4.19 – 4.15 (m, 1H), 4.05 – 3.98 (m, 0.5H), 3.97 – 3.90 (m, 0.5H), 3.88 – 3.80 (m, 0.5H), 3.64 – 3.55 (m, 0.5H), 2.86 – 2.67 (m, 2H), 2.08, 2.06, 2.05 (3s, 9H), 1.97 (s, 3H), 1.45 (m, 9H). ^13^C NMR (150 MHz, CDCl3) δ 80.3 (d, J = 41.8, ^13^C-1), 73.5 (t, J = 42.7, ^13^C-5), 72.8 (t, J = 39.9, ^13^C-3), 67.5 (t, J = 41.8, ^13^C-2), 61.6 (d, J = 44.5, ^13^C-6), 53.5 (t, J = 39.8, ^13^C-2). HRMS (ESI/Orbitrap) m/z [M+H]^+^ Calcd for C_37_H_47_N_3_O_13_^+^: 746.3226; found, 746.3207.

N-*α*-(Fluorenylmethoxycarbonyl)-N-*γ*-(2-acetamido-3,4,6-tri-*O*-acetyl-2-deoxy-*β*-D-glucopyranosyl)-L-asparagine (**4**). A solution of compound **3** (195 mg, 0.26 mmol) in formic acid (5 mL) was stirred overnight at room temperature. At this point, TLC (silica, 5:1 CH_2_Cl_2_–MeOH) showed the reaction was complete. The solvent was evaporated in vacuum and the residue was purified by flash silica gel chromatography (0−15% MeOH in CH_2_Cl_2_) to give **4** (164 mg, 92%) as a pale solid. ^1^H NMR (600 MHz, CDCl_3_) δ 7.76 (d, J = 7.4 Hz, 2H, Fmoc-Ar), 7.59 (d, J = 7.4 Hz, 2H, Fmoc-Ar), 7.40 (t, J = 7.4 Hz, 2H, Fmoc-Ar), 7.49 – 7.42 (m, 1H), 7.30 (t, J = 7.4 Hz, 2H, Fmoc-Ar), 6.35 (d, J = 7.2 Hz, 1H), 6.19 (d, J = 8.0 Hz, 1H), 5.30 – 5.20 (m, 1.5H), 5.05 – 4.95 (m, 1.5H), 4.65 – 4.54 (m, 1H), 4.47 – 4.15 (m, 4H), 4.08 – 3.98 (m, 0.5H), 3.97 – 3.92 (m, 0.5H), 3.91 – 3.85 (m, 0.5H), 3.68 – 3.61 (m, 0.5H), 2.90 – 2.75 (m, 2H), 2.09, 2.07, 2.04, 2.00 (4s, 12H). ^13^C NMR (150 MHz, CDCl_3_) δ 80.3 (d, J = 41.1, ^13^C-1), 73.6 (t, J = 43.2, ^13^C-5), 72.7 (t, J = 39.7, ^13^C-3), 67.7 (t, J = 41.6, ^13^C-2), 61.6 (d, J = 44.3, ^13^C-6), 53.4 (t, J = 39.7, ^13^C-2). HRMS (ESI-TOF) m/z [M+H]^+^ Calcd for C_33_H_38_N_3_O_13_^+^: 690.2600; found, 690.2580.
